# Flexibility in Gene Coexpression at Developmental and Evolutionary Timescales

**DOI:** 10.1101/2024.12.10.627761

**Authors:** Eva K Fischer, Youngseok Song, Wen Zhou, Kim L Hoke

**Author notes:** Author for correspondence: Eva K Fischer Department of Neurobiology, Physiology and Behavior University of California Davis One Shields Avenue Davis, CA 95616.

## Abstract

The explosion of next-generation sequencing technologies has allowed researchers to move from studying single genes to thousands of genes, and thereby to also consider the relationships within gene networks. Like others, we are interested in understanding how developmental and evolutionary forces shape the expression of individual genes, as well as the interactions among genes. In pursuing these questions, we confronted the central challenge that standard approaches fail to control the Type I error and/or have low power in the presence of high dimensionality (i.e., large number of genes) and small sample size, as in many gene expression studies. To overcome these challenges, we used random projection tests and correlation network comparisons to characterize differences in network connectivity and density. We detail central challenges, discuss sample size guidelines, and provide rigorous statistical approaches for exploring coexpression differences with small sample sizes. We apply these approaches in a species known for rapid adaptation – the Trinidadian guppy (*Poecilia reticulata*). Our findings provide evidence for coexpression network differences at developmental and evolutionary timescales and suggest that flexibility in gene coexpression relationships could promote evolvability.

## INTRODUCTION

Genes neither act nor evolve in isolation. Rather, genes are members of physically and functionally interacting networks. There is long-standing interest in how the nature of these interactions influences the extent to which gene sequence and expression changes are constrained at developmental and evolutionary timescales, and thereby influence the evolvability of higher-order phenotypes. On the one hand, genes with many interaction partners (i.e., those that occupy central ‘hub’ positions in a network) may be targets of developmental switches and evolutionary selection because they most strongly influence network output and phenotypic change (Chateigner *et al*., 2020; Friedman *et al*., 2020). Alternatively, expression changes in highly connected genes may be constrained by pleiotropic effects imposed by their many connections (Jeong *et al*., 2001; Hahn & Kern, 2005), and developmental and evolutionary changes may therefore be more prevalent in peripheral genes that have lower connectivity and lower pleiotropic loads (Kim *et al*., 2007; Mähler *et al*., 2017). Despite their contrasting predictions, both hypotheses rely on the assumption that connections among genes – and therefore network topology – are stable. If the relationships among genes are instead flexible across developmental and evolutionary timescales, gene network position may not be a good predictor of evolvability.

The role of interactions among genes in evolutionary processes has historically been difficult to test because physical interaction networks were well-characterized in only very few species (e.g. protein networks in yeast (Jeong *et al*., 2001; Hahn *et al*., 2004; Jovelin & Phillips, 2009)) and simultaneously surveying expression in large numbers of genes was challenging, if not impossible. The proliferation of next generation sequencing technologies, and specifically RNA-sequencing (hereafter RNAseq), has removed these constraints from a technical perspective. Although network and coexpression analyses of RNAseq datasets remain less common than gene-wise differential expression analyses, studies in this area provide intriguing – albeit conflicting – results. In support of the idea that genes in central network positions are evolutionary constrained, while those in the network periphery are more evolvable, are studies demonstrating that centrality in coexpression networks is negatively correlated with divergence in gene expression (Warnefors & Kaessmann, 2013; Mähler *et al*., 2017; Kuo *et al*., 2023) and sequence evolution (Josephs *et al*., 2017; Masalia *et al*., 2017; Harnqvist, 2021), while genes in peripheral positions show greater magnitude expression divergence (Mähler *et al*., 2017) and signatures of positive selection (Kim *et al*., 2007; Mähler *et al*., 2017). In contrast, other studies provide evidence for a bias toward changes in the expression of and selection on genes with high centrality in coexpression networks (Koubkova-Yu *et al*., 2018; Chateigner *et al*., 2020; Friedman *et al*., 2020; Rennison & Peichel, 2022), supporting the alternative hypothesis that central genes better predict phenotypic variation and are therefore targets of selection. Attempts to resolve these conflicts include hypotheses that preferential divergence in central versus peripheral genes may be associated with different types of traits (Warnefors & Kaessmann, 2013; Des Marais *et al*., 2017) or distinct selective modes and timescales (Luisi *et al*., 2015), or that intermediate levels of pleiotropy facilitate evolution (Hämälä *et al*., 2020). These ideas are intriguing, yet also rely on the assumption that connections among genes are stable rather than flexible.

If the relationships among genes are instead flexible, then the degree of pleiotropy and its presumed consequences are not fixed, and flexibility in coexpression relationships among genes could reduce pleiotropic load (e.g., (Wang et al., 2010;Pavlicev & Wagner, 2012; Pavličev & Cheverud, 2015). In other words, individual genes may be more able to change in expression and drive phenotypic change if their interactions with other genes can be altered to minimize off-target effects. Conversely, flexibility in coexpression relationships might improve the ability of underlying gene expression networks to buffer higher-level phenotypes through homeostatic change (e.g., (Fischer *et al*., 2016; Badyaev, 2018; Hoke *et al*., 2019)). Importantly, either scenario implies that the relationships among genes may themselves be targets of selection. Alternatively, changes in gene coexpression could represent drift and/or transcriptional noise if these changes do not amount to selectable differences at the network and/or organismal level. While this last scenario is less interesting from an adaptationist perspective, such ‘neutral’ changes may nonetheless have consequences for evolutionary trajectories, for example by giving rise to cryptic variation that is revealed under novel environmental conditions (West-Eberhard, 2003; McGuigan & Sgrò, 2009; Paaby & Rockman, 2014). In brief, all the above alternatives highlight that coexpression relationships could vary at developmental and evolutionary timescales with consequences in the short- and long-term.

Interest in coexpression analyses is being met by a growing collection of tools for (differential) gene coexpression analysis (Wang *et al*., 2017; Chowdhury *et al*., 2020; Tommasini & Fogel, 2023). While these software packages make advanced network analyses accessible, they do not eliminate the statistical limitations of these approaches. These limitations arise primarily from the combination of small sample sizes and high-dimensional data (tens of thousands of genes) emblematic of transcriptomic studies. Sample sizes have increased as sequencing costs have decreased, yet per group sample sizes commonly remain less than ten. Pooling samples across experimental groups or multiple studies can bring the total experimental sample size into the range recommended for network analyses (e.g., N=20 by (Langfelder & Horvath, 2008; Ballouz *et al*., 2015)); however, this solution is not viable when the goal is to test if experimental groups differ in coexpression structure. Moreover, correlations in expression derived from experimental groups that differ in their underlying transcriptional networks are particularly hard to interpret. This leaves researchers trapped between an experimental rock and hard place.

Like others, we are interested in using transcriptomic analyses to understand the biological basis of complex phenotypes, and specifically in exploring changes in individual genes as well as the interactions among genes. To this end, we characterized the effects of genetic background (high-predation versus low-predation populations) and developmental environment (rearing with and without predator chemical cues) on brain gene coexpression in two parallel, independent evolutionary lineages of Trinidadian guppies (*Poecilia reticulata*). In Trinidad, downstream, high-predation fish have repeatedly and independently colonized upstream, low-predation environments (Gilliam *et al*., 1993; Barson *et al*., 2009F; Willing *et al*., 2010a; Fraser *et al*., 2015), leading to parallel adaptive changes in life-history, morphology, and behavior (Reznick *et al*., 1990, 2001; Endler, 1995; Reznick, 1997; Magurran, 2005a). In other words, each river drainage represents a naturally replicated experiment demonstrating parallel phenotypic adaptation.

Prior work has demonstrated that both evolutionary history with predators and developmental experience with predators shape life history (Torres Dowdall *et al*., 2012), morphology (Torres-Dowdal *et al*., 2012; Fischer *et al*., 2013; Ruell *et al*., 2013; Handelsman *et al*., 2014), physiology (Handelsman *et al*., 2013; Fischer *et al*., 2014), and behavior (Huizinga *et al*., 2009; Torres-Dowdall *et al*., 2012; Fischer *et al*., 2016b). Here, we extend a prior analysis (Fischer et a. 2021) that characterized developmental plasticity and genetic differences in transcriptome-wide expression levels using animals from different populations reared with and without exposure to predator cues (Figure 1). To address plasticity and divergence in the coexpression patterns in these same data, we first asked whether connectivity patterns among genes differed based on evolved differences between lineages (i.e., river drainages), adaptive differences between high- and low-predation populations of the same lineage, and developmental differences arising from rearing with or without predators. In essence, these comparisons allowed us to test whether changes in gene coexpression are a feature of divergence across timescales (genetic and developmental) and parallel, independent evolutionary events. Following these comparisons, we further investigated whether connectivity influenced a gene’s propensity for expression divergence.

**Figure 1.**
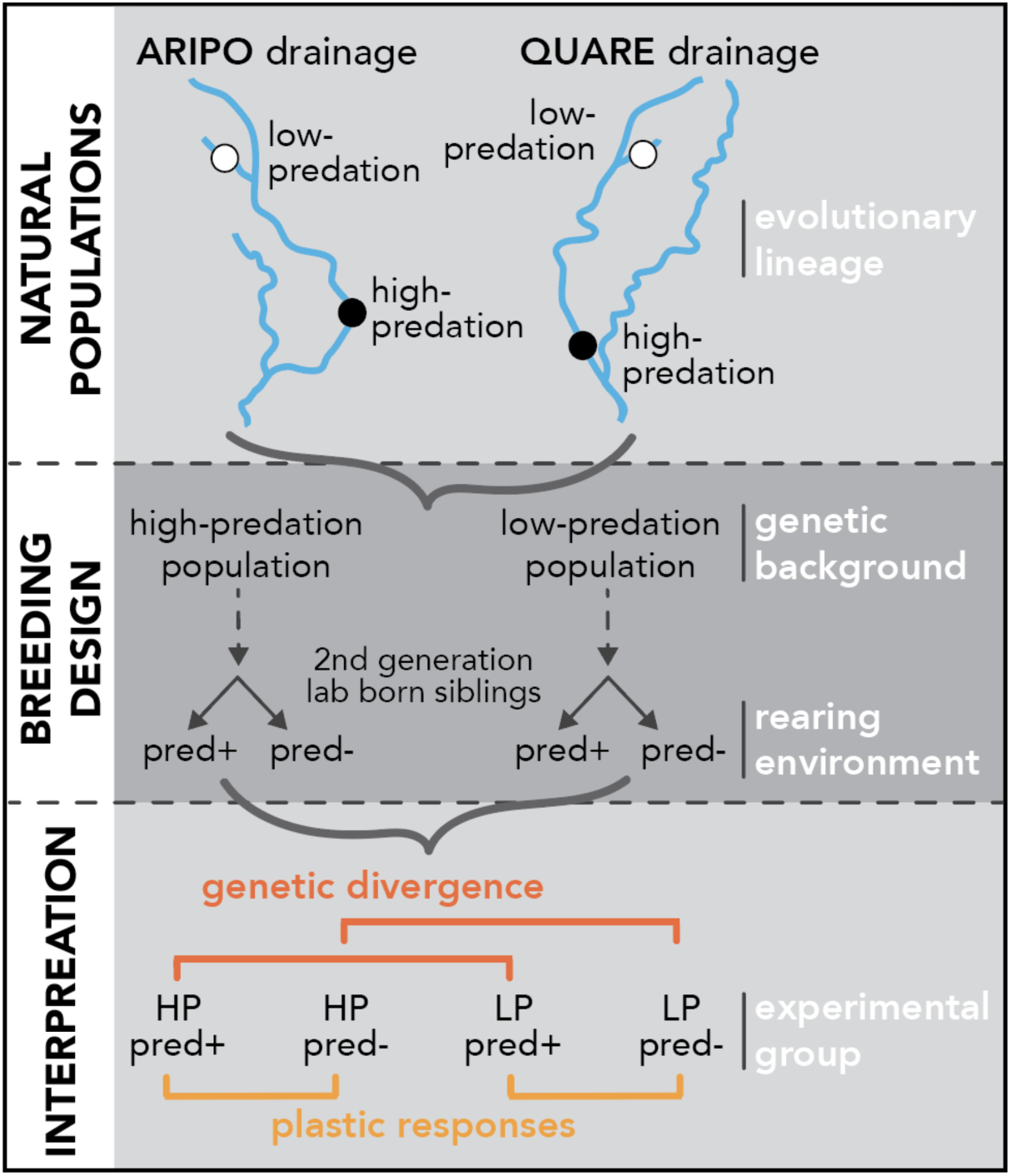
Conceptual overview and interpretation. Top: Natural populations in different river drainages represent distinct evolutionary lineages. Middle: Overview of laboratory breeding design disentangling genetic and environmental effects. Bottom: Interpretation of comparisons of interest between experimental groups resulting from 2×2 breeding design shown above. HP = high-predation, LP = low-predation, pred+ = reared with predator chemical cues, pred- = reared without predator chemical cues. Modified from Fischer et al. 2021.

In pursuing these questions, we confronted the statistical challenges that standard approaches may fail to control Type I error and/or have low power when dimensionality is high (i.e., large number of genes) and sample size is small. From a technical perspective, we present a case study for those interested in (differential) coexpression with small per-group sample sizes. We discuss key challenges, set clear sample size guidelines to control Type I error while maintaining power, and provide rigorous statistical approaches for exploring coexpression differences even with small sample sizes that can be readily implemented for similar coexpression analyses in other studies. These methods do not replace other, popular approaches but rather complement them to ensure analysis outcomes can be rigorously interpreted. Using these statistical methods, we find evidence for differences in coexpression relationships based on both genetic background and rearing environment, suggesting that changes in the interactions among genes are associated with phenotypic divergence at developmental and evolutionary timescales.

## METHODS

### Fish collection and rearing

Samples here are the same as those described in (Fischer *et al*., 2021). Guppies used in this study were second generation lab-born fish from unique family lines established from wild-caught high-predation (HP) and low-predation (LP) populations in the Aripo and Quare river drainages in the Northern Range mountains of Trinidad. At birth, we split second-generation siblings into rearing environments with (pred+) or without (pred-) predator chemical cues, and they remained in these environments until the completion of the experiment (Figure 1) (as in (Fischer *et al*., 2016b)). Guppies were individually housed in 12:12 hour light cycle and fed a measured food diet once daily. All experimental methods were approved by the Colorado State University Animal Care and Use Committee (Approval #12-3818A).

In brief, each drainage consists of a 2×2 factorial design that distinguishes genetic from developmental effects of predation (Figure 1). Pair-wise comparisons of biological relevance are: (1) HP pred-vs LP pred-, an experiment comparing populations reared in an environment lacking predator cues to identify genetic differences between populations; (2) HP pred+ vs HP pred-, to identify environmentally induced changes mimicking the situation in which high-predation fish colonize low-predation environments, i.e., “ancestral plasticity”; (3) LP pred-vs LP pred+, to identify environmentally induced changes comparable to the situation in which low-predation fish are washed downstream and a measure of whether ancestral plasticity is maintained in the derived population; (4) HP pred+ vs LP pred+, to identify genetic differences when fish are raised with environmental cues of predation. We also compared the same experimental groups across drainages (e.g. HP pred+ in Aripo drainage vs HP pred+ in Quare drainage) to understand differences associated with parallel, independent evolutionary lineages.

### Tissue collection and processing

We collected brain tissue from mature males in the groups described above 10 minutes after lights on in the morning. We extracted RNA from whole brains using the Qiagen RNeasy Lipid Tissue Mini Kit (Qiagen, Germany) and constructed a sequencing library for each individual using the NEBNext Ultra RNA Library Prep Kit for Illumina (New England Biolabs, Massachusetts, USA). For the Aripo dataset, 40 samples (N=10 per group) were pooled with unique barcodes into eight samples per sequencing library and each library was sequenced on a single lane. For the Quare dataset, 60 samples (N=12-16 per group) were pooled into three sequencing libraries with 20 samples per pool and each library was sequenced in two separate lanes. Libraries were sequenced as 100bp paired-end reads on an Illumina HiSeq 2000 at the Florida State University College of Medicine Translational Science Laboratory (Tallahassee, Florida) in May 2014 (Aripo dataset) and January 2016 (Quare dataset).

### Differential expression analysis

We reported results of differential expression analyses in another study (Fischer *et al*., 2021). We use normalized values and the resulting differential expression status (differentially expressed (DE) versus non differentially expressed (NDE)) as criterion in analyses performed here (see below). Briefly, we normalized read counts using DESeq2 (Love *et al*., 2014) and performed differential expression analysis using the lme4 package in R (github.com/lme4). We used generalized linear mixed models with a negative binomial link to accommodate our experimental design and data type. We included family and sampling week as random effects to identify DE genes for the fixed effects of population of origin (HP / LP), rearing environment (pred+/ pred-), and their interaction. To label DE and NDE genes, we adjusted p-values for multiple hypothesis testing using a direct approach for FDR control (Storey, 2002) as implemented in the fdrtool package in R (Strimmer, 2008). We considered transcripts differentially expressed (DE) if the adjusted p-value was <0.05, and all other genes non differentially expressed (NDE).

Our previous work reported brain gene expression differences based on genetic and developmental influences in both drainages (Fischer *et al*., 2021). We found that genes exhibiting expression changes in response to rearing differences were also more likely to be differentially expressed between high- and low-predation populations. While this pattern was evident in both river drainages, sets of differentially expressed genes were largely non-overlapping between lineages. For two-sample covariance tests and network comparisons (see below), we categorized genes as DE or NDE based on whether they were differentially expressed for the effect of population of origin within each drainage.

### Preliminary analyses

Our initial approach was to characterize and explore gene coexpression using the popular Weighted Gene Correlation Network Analysis (WGCNA) package in R (Langfelder & Horvath, 2008). We calculated module preservation scores using the methods implements in WGCNA which combine a number of difference preservation statistics to calculate a summary preservation score (Langfelder *et al*., 2011). We found that, in both drainages, ∼50% of gene coexpression modules were not preserved across experimental groups (Supplemental Materials). In other words, we identified substantial differences in network structure between groups, suggesting that the common practice of reconstructing coexpression networks by pooling samples across groups may not be valid. In brief, our preliminary analyses using WGCNA underscored the need for a statistical method to discern network differences across groups when dealing with small group sizes (N=10-12 in our case). We sought to address these issues through the alternative statistical approaches detailed below, and further in the Methods and Supplemental Materials. Our approaches are not intended as a replacement for WGCNA and other similar packages but can be used in conjunction to ensure that WGCNA outputs can be rigorously interpreted.

### Challenges from small sample, high dimensional data

In wanting to understand differences in covariance structure and changes in network architecture between our experimental groups, we faced two fundamental challenges. First, controlling Type I error becomes difficult when comparing large networks or covariance structures with limited sample sizes, often leading to spurious discoveries and unreliable results. Also, detecting true effects for separating complicated networks or covariance structures in ultra-high dimensional datasets is difficult due to low statistical power; a common phenomenon in coexpression analysis of RNAseq studies. While there are commonly accepted methods for two sample covariance tests, they either fail to control the Type I error rate with extremely small sample size (e.g., N<25 per group) or have substantially low power (see Supplemental Materials for simulations) (Li & Chen, 2012; Cai *et al*., 2013; Chang *et al*., 2017; Yu *et al*., 2020). Second, comparison of multiple networks is nontrivial, particularly when the networks are of different sizes or have unmatching nodes (Tang *et al*., 2017; Agterberg *et al*., 2020; Qi *et al*., 2024). For gene coexpression analysis these issues apply, for example, when sample sizes vary between groups and are small due to the limitation of experiment constraints, or when comparing subsets of genes of interest that vary in size (e.g., comparing DE vs NDE gene sets, or differently sized coexpression modules) (Agterberg *et al*., 2020; Alyakin *et al*., 2024; Jin *et al*., 2024; Qi *et al*., 2024).

To highlight these challenges, we first conducted simulation studies using existing high-dimensional methods designed for valid inference on comparing large covariance structures with controlled Type I error rate and reasonable power (Li & Chen, 2012; Cai *et al*., 2013; Chang *et al*., 2017; Yu *et al*., 2020), before implementing our method and examining the real data set (see below). We summarize the outcomes of the simulation studies and exploratory comparisons here and refer the interested reader to additional details provided in the Supplemental Materials.

We found that Type I error rates for existing tests were uncontrolled for small sample sizes of N<50 per group, even when the number of genes was relatively small (250 genes, orders of magnitude smaller than what is typical for RNAseq analysis) (Figure S1). In addition to uncontrolled Type I error, the existing methods were substantially underpowered for small sample sizes. Specifically, the empirical power was overall low (<0.25) for sample sizes N<30 per group (Figure S2). These issues plagued our dataset, which is representative of most RNAseq studies exploring connections between gene expression and behavior (per group samples N<10, ∼20,000+ genes). Importantly, these issues are not resolved by subsampling the data to include a smaller number of genes (Figure S3), an approach commonly deployed by network analysis packages (e.g., filtering for the 5,000-8,000 most variable genes in WGCNA or the approach of (Qiu *et al*., 2021)).

### Our new high-dimensional covariance comparison

The task of determining whether a common covariance structure can be assumed between two groups for downstream analysis can be addressed using two-sample covariance tests. If this assumption is supported by the data, pooling the samples across groups can improve statistical efficiency in recovering the underlying network and provide a better understanding of its structure. However, from the above simulations, it was clear that existing approaches to compare large covariance structures fail even when using only a small subset of genes. To overcome these issues, we extended the random projection-based covariance test (Wu & Li, 2020) to develop a new two-sample comparison method suitable for our data (see Results and Supplemental Materials).

We conduct our newly developed random projection-based tests on residuals from the linear mixed model described above and in (Fischer *et al*., 2021). We used residuals to remove the mean effects of population of origin and the rearing environment, thereby allowing us to focus on the underlying co-expression patterns among genes, rather than differences in mean expression, when studying dependency structures among genes. We focused on pairwise comparisons of biological interest (Figure 1c and above). Within each drainage, we compared: (1) HP pred-vs LP pred-, (2) HP pred+ vs HP pred-, (3) LP pred-vs LP pred+, and (4) HP pred+ vs LP pred+. We also compared the same experimental groups across drainages (e.g. HP pred+ in Aripo drainage vs HP pred+ in Quare drainage) to understand differences associated with parallel, independent evolutionary lineages. For both within and between drainage comparison, we considered the four comparisons jointly to control family-wise error rate.

### Correlation network comparisons

We used correlation network analyses to characterize differences in network structure that accompany the changes in covariance matrices. Because differential expression might drive patterns of differential coexpression patterns, we tested whether DE versus NDE genes differed in their connectivity with other genes and analyzed genetic and developmental influences on network structure separately in DE and NDE gene sets. We used the DE and NDE characterizations determined previously using generalized linear models to detect population differences as summarized above, although we note that the sets of differentially expressed genes were largely non-overlapping between evolutionary lineages (drainages). A challenge with this analysis was the lack of consensus to define networks, in addition to the fact that derived networks will usually have very different sizes and unmatched nodes (i.e., NDE genes far outnumber DE genes, a single gene is inherently in only one category, and the DE genes in the Quare drainage outnumbered those in Aripo).

The network comparisons involved two steps: (i) reconstruction of the coexpression network using gene-wise correlations, and (ii) comparing two networks of different sizes. To achieve (i), we first tested whether the correlation between each pair of genes was zero, while controlling the FDR using the method from (Cai & Liu, 2016), as detailed in Supplemental Materials B4.3. Using this method, an undirected edge is drawn between any two genes (nodes) with nonzero correlation, forming the coexpression network. To assess the constructed network’s sensitivity to different FDR levels, we compared network summary plots at multiple FDR cutoffs (α = 0.01, 0.05, 0.1). If two networks are distinct, their summary plots will differ (Maugis *et al*., 2017). The network summary plots (Figure S6) suggest that the correlation-based coexpression network is relatively insensitive to different FDR levels. Therefore, we used a coexpression network with α = 0.05 for all subsequent analyses

We compared network pairs of interest to examine their structural differences, using a network summary plot and two-sample network tests based on relative frequencies of different subgraphs adjusted for the sparsity (density of edges) of networks (Maugis et al., 2017; Shao et al., 2022). A major concern in comparing DE versus NDE networks, which has been largely overlooked in literature, is that the different collections of genes in these two sets (e.g., DE genes are a small subset of all genes) make the two corresponding networks have different numbers of unmatchable nodes (Qi *et al*., 2024). To address this, we adopted the network comparison test proposed by (Shao *et al*., 2022), which accommodates networks of different sizes by analyzing their network moments for specific motifs using the difference of two subgraph densities adjusted for their edge densities (additional details in Supplemental Materials). Comparing subgraph densities provides a measure of network topology and sparsity that can be visualized using network summary plots. To help readers interpret plots of the real data, we provide a set of example networks and their corresponding network summary plots (Figure 2 and Glossary).

**Figure 2.**
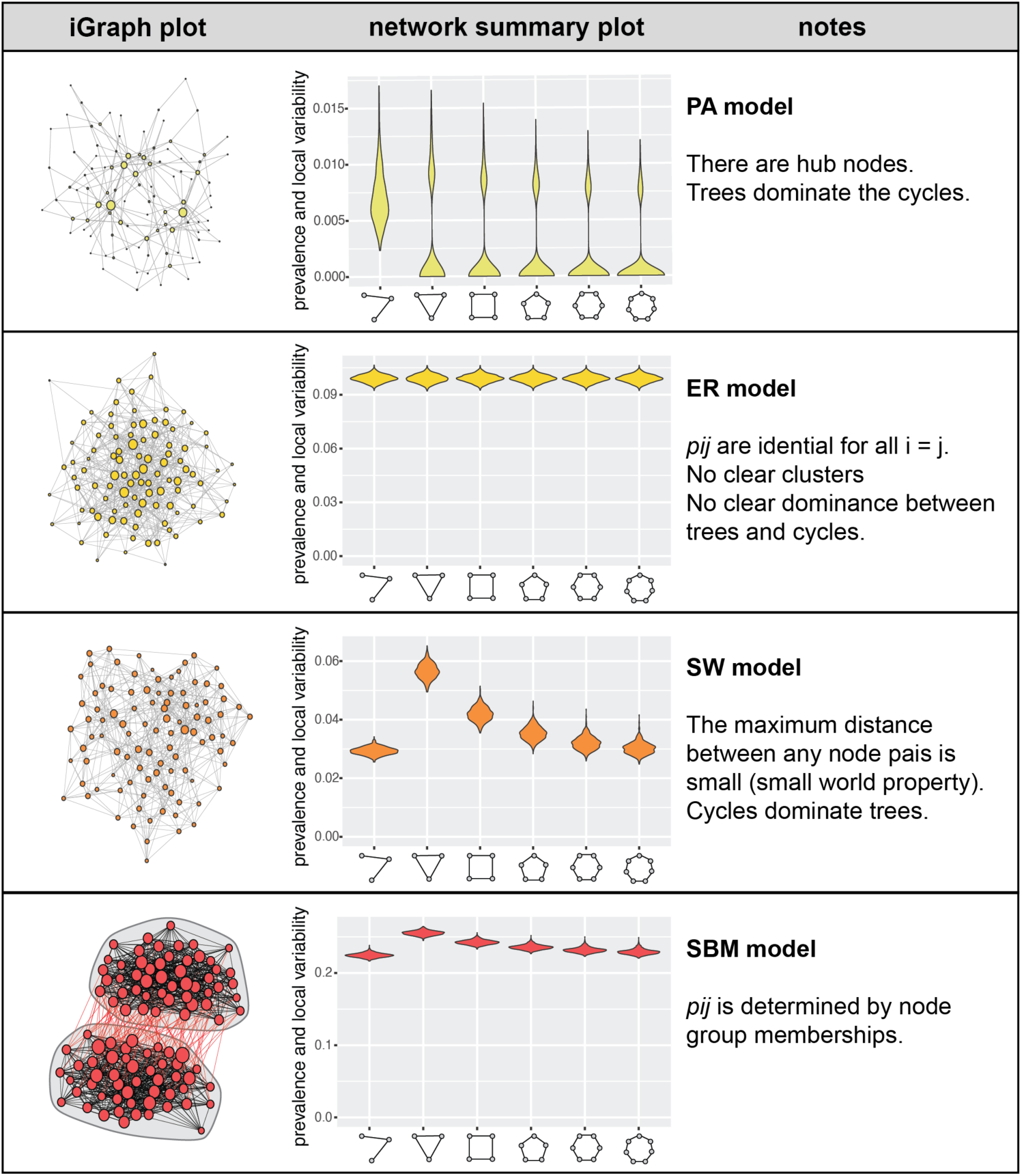
Example network summary plots. Sample networks (iGraphs), their corresponding network summary plots, and a brief interpretation are shown. Examples are intended to help readers unfamiliar with network summary plots interpret summary plots of the real data.

We made various comparisons of the different subgraphs to understand differences based on rearing environment, population of origin, and evolutionary lineage. We conducted one-sided comparisons for each of the three sub-graph motifs (v-shape, triangle, and 3-star). Comparisons fell into four general categories: (1) comparing DE and NDE gene sets within each experimental group, (2) comparing developmental differences (i.e., rearing with (pred+) or without (pred-) predators) for DE and NDE gene sets, (3) comparing populations differences (i.e., high-predation versus low-predation) for DE and NDE gene sets, and (4) comparing experimental groups across evolutionary lineages (i.e., Aripo vs Quare drainage) for DE and NDE gene sets.

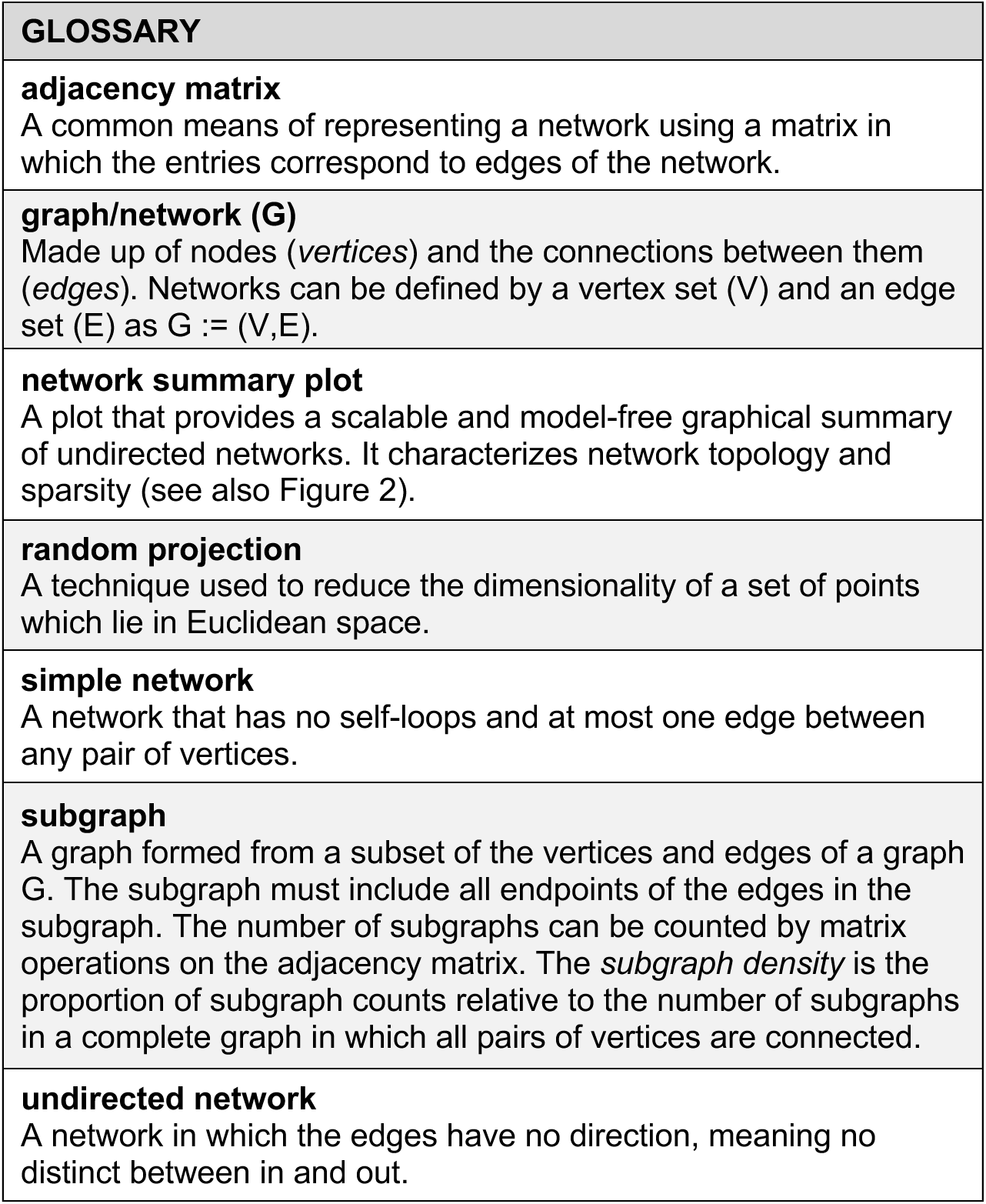

| GLOSSARY |
| --- |
| **adjacency matrix**<br>A common means of representing a network using a matrix in which the entries correspond to edges of the network. |
| **graph/network (G)**<br>Made up of nodes (*vertices*) and the connections between them (*edges*). Networks can be defined by a vertex set (V) and an edge set (E) as $G := (V, E)$. |
| **network summary plot**<br>A plot that provides a scalable and model-free graphical summary of undirected networks. It characterizes network topology and sparsity (see also Figure 2). |
| **random projection**<br>A technique used to reduce the dimensionality of a set of points which lie in Euclidean space. |
| **simple network**<br>A network that has no self-loops and at most one edge between any pair of vertices. |
| **subgraph**<br>A graph formed from a subset of the vertices and edges of a graph G. The subgraph must include all endpoints of the edges in the subgraph. The number of subgraphs can be counted by matrix operations on the adjacency matrix. The *subgraph density* is the proportion of subgraph counts relative to the number of subgraphs in a complete graph in which all pairs of vertices are connected. |
| **undirected network**<br>A network in which the edges have no direction, meaning no distinct between in and out. |

## RESULTS

### Changes in coexpression networks based on genetics and environment

We were interested in comparing gene coexpression patterns based on genetic background and rearing environment. To overcome problems associated with limited sample sizes yet high-dimensional data, we used random projection-based tests to compare covariance structures between experimental groups (Wu & Li, 2020). Instead of testing two large covariance matrices directly, these tests compute random projections of the data into lower dimensional spaces and test the equality of variances. Previous work (Wu & Li, 2020) deployed this approach in one-dimensional space, and we developed a generalized version for multi-dimensional space (further details in Supplemental Materials). Using simulations, we confirmed this approach controls type-I error (Figure S4) rate while maintaining power (Figure S5).

Applying our method to the real data, we considered the set of all genes that passed filtering criteria (Aripo: 13,446; Quare: 14,379). We found significant differences in covariance structures between high-predation and low-predation fish reared with predators (HP pred+ vs LP pred+) in both drainages (Figure 3 Table 1). Analysis of the Quare dataset found a marginally significant difference between high-predation fish reared with and without predators (HP pred+ vs HP pred-). We also compared the covariance structures between the same treatment groups across drainages. Here, we found significant differences in all comparisons (Table 2). In short, when considering all genes in the daataset, we found evidence for changes in gene coexpression based on evolutionary lineage (drainage), genetic background (population), and rearing environment.

**Figure 3.**
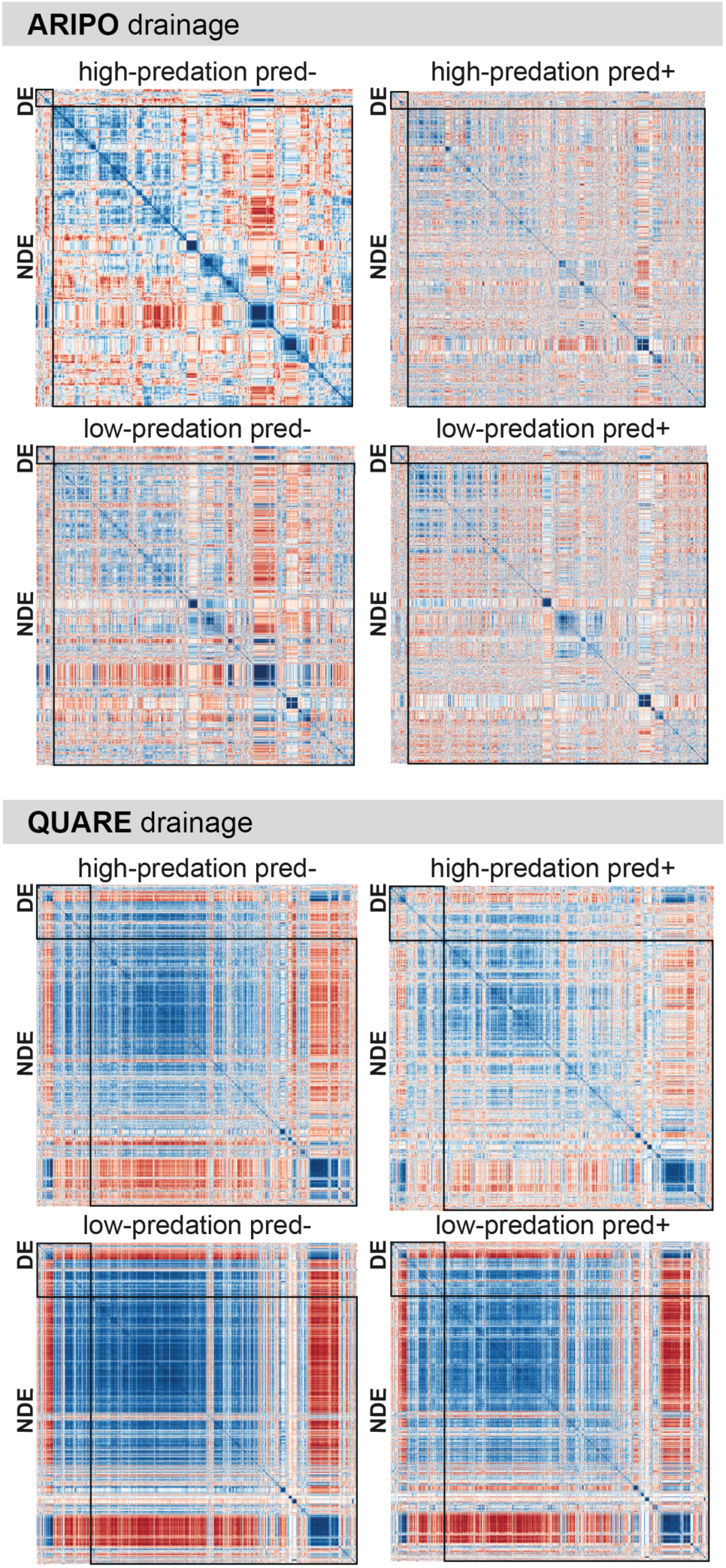
Visualization of coexpression differences between experimental groups and evolutionary lineages. Heatmaps provide a visualization of statistical differences in Pearon’s correlations of expression between genes based on genetic background (high-vs low-predation), rearing environment (with (pred+) or without (pred-) predators), and evolutionary lineage (Aripo and Quare drainage). Gene order is determined by hierarchical clustering of the high-predation pred-group, meaning that the same position in two heatmaps represents the correlation of identical pairs of genes. For ease of visualization and computation, only the 1,000 most variable genes are shown.

**Table 1.**
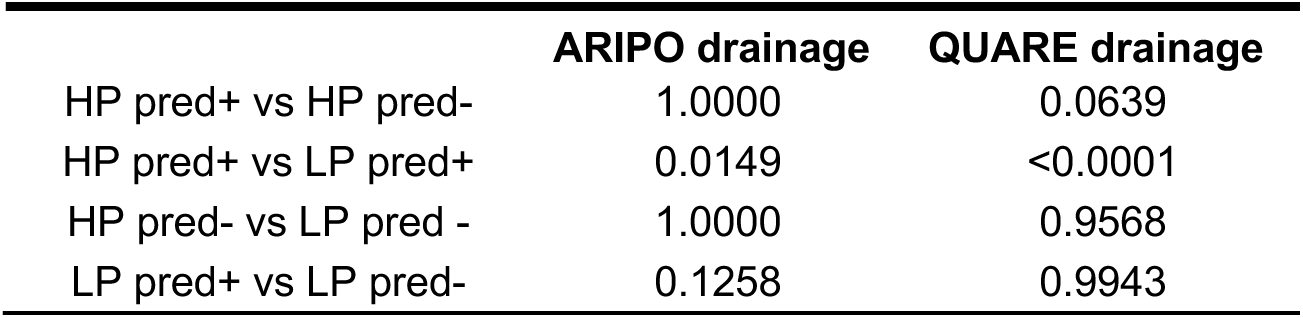
Approximated p-values from random projection tests comparing covariance structure for all genes (DE and NDE) between treatment groups.

**Table 2.** Approximated p-values from random projection tests comparing covariance structure across drainages.

| **ARIPO vs QUARE** | **p-value** |
| --- | --- |
| HP pred- | <0.0001 |
| HP pred+ | 0.0001 |
| LP pred - | 0.0001 |
| LP pred+ | 0.0107 |

### Coexpression networks and differential expression

To further characterize the changes in coexpression among genes, we compared the prevalence of network motifs across treatment groups. Because we reasoned that divergence in expression levels might itself impact networks, we first addressed whether coexpression networks differed between differentially expressed (DE) and non-differentially expressed (NDE) genes. We performed these comparisons separately for each treatment group, given group-level differences in covariance structures detailed above. Further, because our networks include different gene sets of different sizes, we adopted network comparisons of subgraph densities adjusted for overall edge density (Shao *et al*., 2022).

We compared the v-shape (subgraphs with three nodes and two edges), triangle (subgraphs with three nodes and three edges), and 3-star (subgraphs with four nodes and three edges) prevalence within networks relative the amounts expected given edge densities (see visualizations in Figures 2, 4, and S9). We considered these specific subgraphs commonly used in the network literature as metrics of connectivity and clusterability (Tang *et al*., 2017; Agterberg *et al*., 2020; Qi *et al*., 2024) because all more complex motifs can be deconstructed to these components, and this set of motifs is therefore sufficient to capture network differences such that a greater abundance of complex motifs indicates higher network connectivity and density. Finding different patterns in DE and NDE gene sets within treatment groups (Figure 4, Table S10), we compared network motifs among treatment groups separately in the DE and NDE gene sets. Overall, we found differences in the prevalence of v-shape, triangle, and 3-star motifs in coexpression networks based on genetic and environmental influences, but without a simple rule of directionality (Figure 4, Tables 3 and 4).

**Figure 4.**
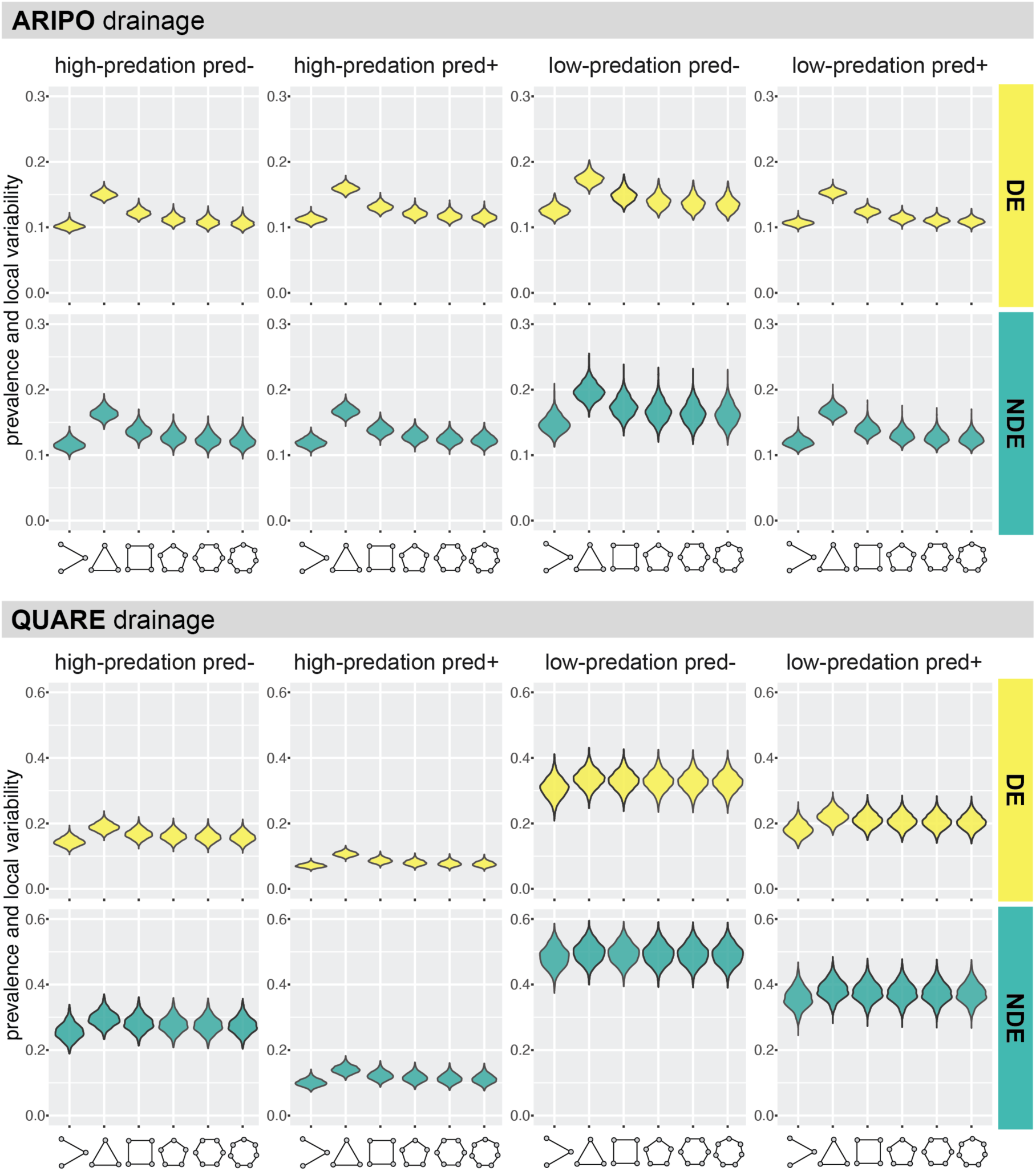
Network summary plots of correlations networks. Plots provide a visualization of network topology and sparsity for networks of DE (yellow) and NDE (teal) genes that differed in mean expression levels between the two populations in each river drainage. Subgraph shapes are depicted on the x-axis. See Tables 3 and 4 for outcomes of statistical comparisons between groups.

**Table 3.**
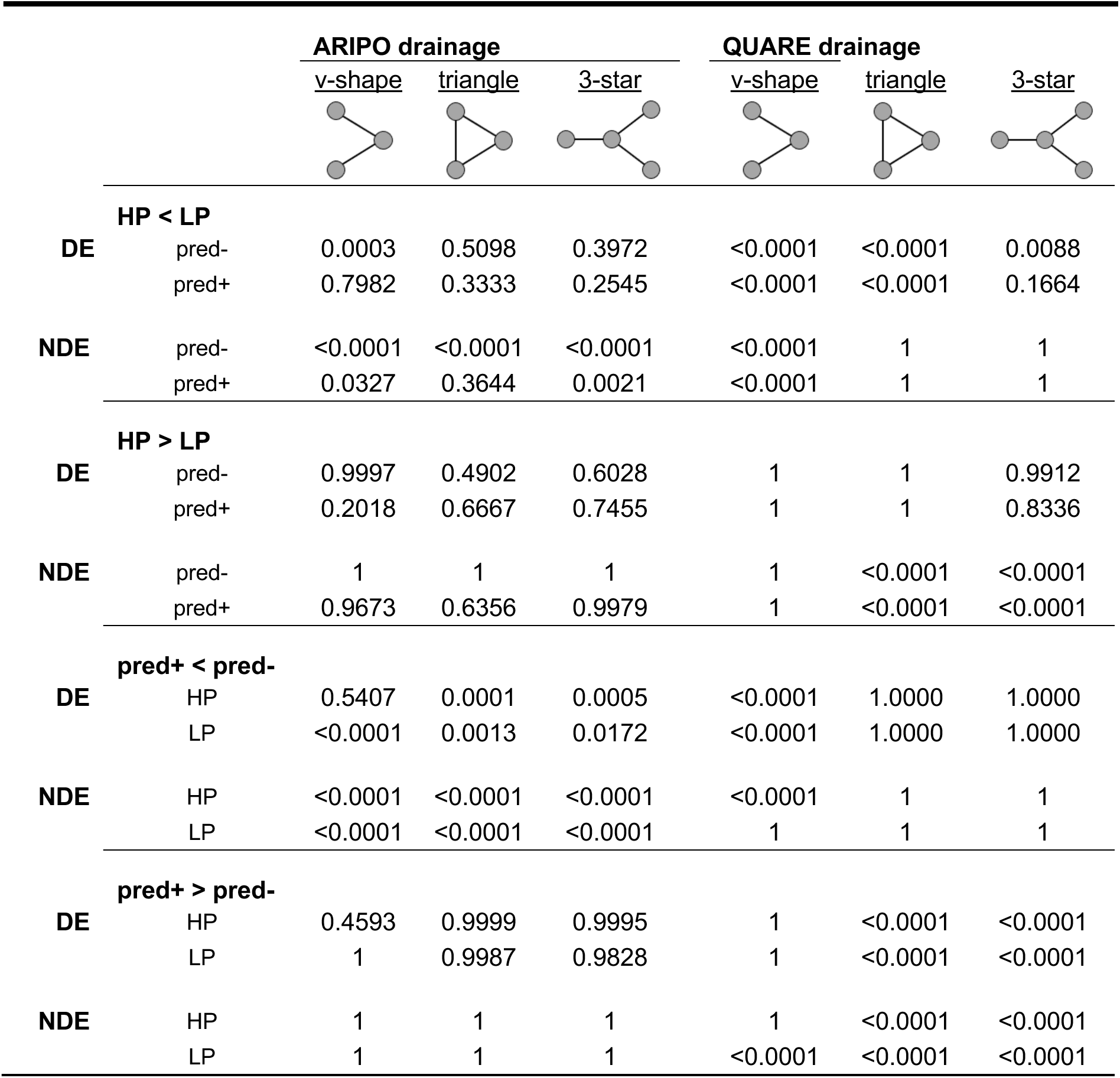
Network comparisons of treatment groups for DE and NDE genes across datasets. Comparisons of sparsity-adjusted subgraph densities tested the alternatives that gene networks had smaller or larger subgraph density than other networks based on differences in population of origin and rearing environment. P-values from one-sided alternative tests are reported for the v-shape, triangle, and 3-star subgraph types.

|  |  | **ARIPO drainage** |  |  | **QUARE drainage** |  |  |
| --- | --- | --- | --- | --- | --- | --- | --- |
|  |  | <u>v-shape</u> | <u>triangle</u> | <u>3-star</u> | <u>v-shape</u> | <u>triangle</u> | <u>3-star</u> |
| **HP < LP** |  |  |  |  |  |  |  |
| **DE** | pred- | 0.0003 | 0.5098 | 0.3972 | <0.0001 | <0.0001 | 0.0088 |
|  | pred+ | 0.7982 | 0.3333 | 0.2545 | <0.0001 | <0.0001 | 0.1664 |
| **NDE** | pred- | <0.0001 | <0.0001 | <0.0001 | <0.0001 | 1 | 1 |
|  | pred+ | 0.0327 | 0.3644 | 0.0021 | <0.0001 | 1 | 1 |
| **HP > LP** |  |  |  |  |  |  |  |
| **DE** | pred- | 0.9997 | 0.4902 | 0.6028 | 1 | 1 | 0.9912 |
|  | pred+ | 0.2018 | 0.6667 | 0.7455 | 1 | 1 | 0.8336 |
| **NDE** | pred- | 1 | 1 | 1 | 1 | <0.0001 | <0.0001 |
|  | pred+ | 0.9673 | 0.6356 | 0.9979 | 1 | <0.0001 | <0.0001 |
| **pred+ < pred-** |  |  |  |  |  |  |  |
| **DE** | HP | 0.5407 | 0.0001 | 0.0005 | <0.0001 | 1.0000 | 1.0000 |
|  | LP | <0.0001 | 0.0013 | 0.0172 | <0.0001 | 1.0000 | 1.0000 |
| **NDE** | HP | <0.0001 | <0.0001 | <0.0001 | <0.0001 | 1 | 1 |
|  | LP | <0.0001 | <0.0001 | <0.0001 | 1 | 1 | 1 |
| **pred+ > pred-** |  |  |  |  |  |  |  |
| **DE** | HP | 0.4593 | 0.9999 | 0.9995 | 1 | <0.0001 | <0.0001 |
|  | LP | 1 | 0.9987 | 0.9828 | 1 | <0.0001 | <0.0001 |
| **NDE** | HP | 1 | 1 | 1 | 1 | <0.0001 | <0.0001 |
|  | LP | 1 | 1 | 1 | <0.0001 | <0.0001 | <0.0001 |

**Table 4.**
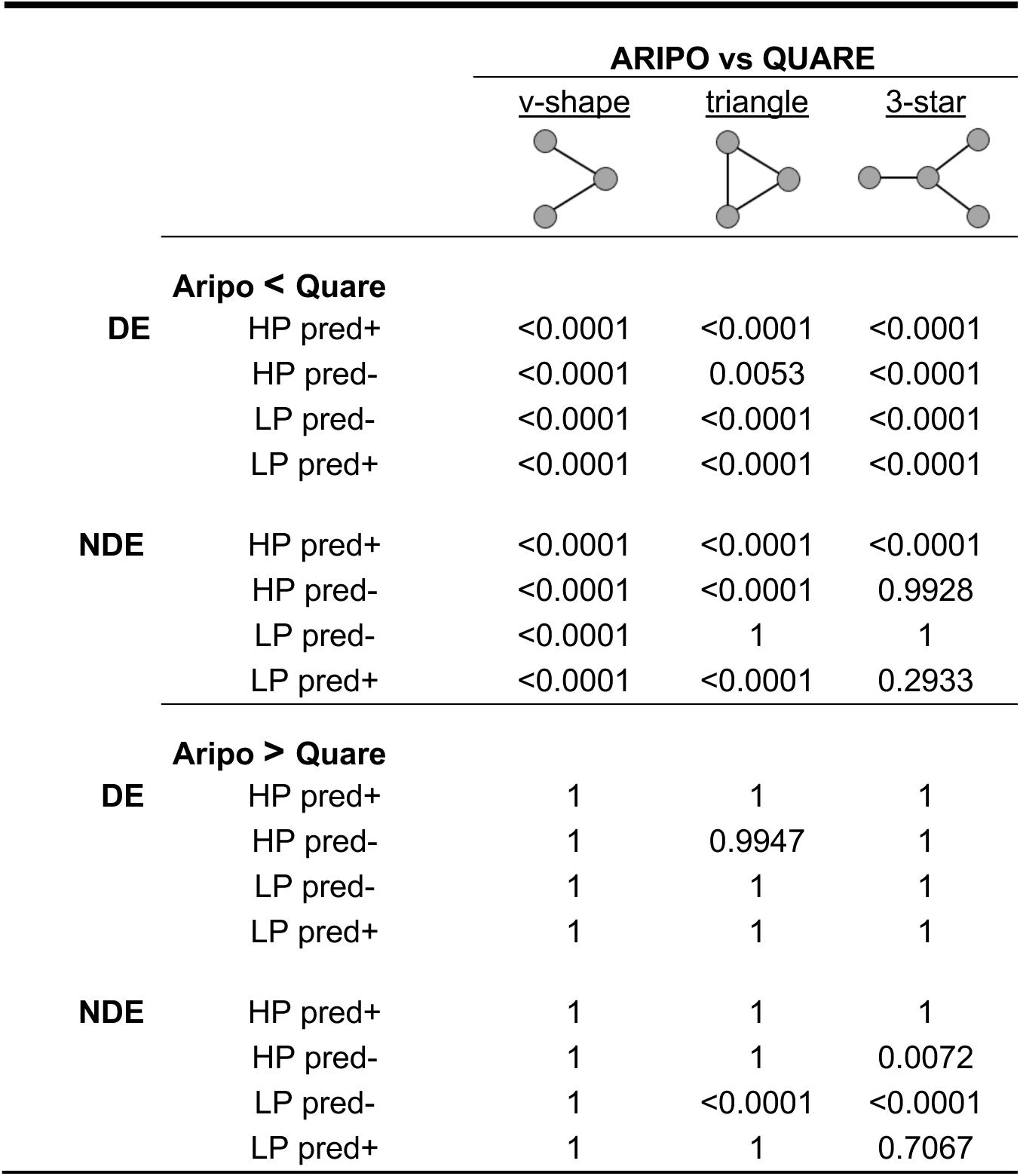
Network comparisons of datasets within treatment groups for DE and NDE genes. Comparisons of sparsity-adjusted subgraph densities tested the alternatives that gene networks for each treatment group in one drainage had smaller or larger subgraph density than networks for the same treatment group in the other drinage. P-values from one-sided alternative tests are reported for the v-shape, triangle, and 3-star subgraph types.

|  |  | **ARIPO vs QUARE** |  |  |
| --- | --- | --- | --- | --- |
|  |  | <u>v-shape</u> | <u>triangle</u> | <u>3-star</u> |
| **Aripo < Quare** |  |  |  |  |
| **DE** | HP pred+ | <0.0001 | <0.0001 | <0.0001 |
|  | HP pred- | <0.0001 | 0.0053 | <0.0001 |
|  | LP pred- | <0.0001 | <0.0001 | <0.0001 |
|  | LP pred+ | <0.0001 | <0.0001 | <0.0001 |
| **NDE** | HP pred+ | <0.0001 | <0.0001 | <0.0001 |
|  | HP pred- | <0.0001 | <0.0001 | 0.9928 |
|  | LP pred- | <0.0001 | 1 | 1 |
|  | LP pred+ | <0.0001 | <0.0001 | 0.2933 |
| **Aripo > Quare** |  |  |  |  |
| **DE** | HP pred+ | 1 | 1 | 1 |
|  | HP pred- | 1 | 0.9947 | 1 |
|  | LP pred- | 1 | 1 | 1 |
|  | LP pred+ | 1 | 1 | 1 |
| **NDE** | HP pred+ | 1 | 1 | 1 |
|  | HP pred- | 1 | 1 | 0.0072 |
|  | LP pred- | 1 | <0.0001 | <0.0001 |
|  | LP pred+ | 1 | 1 | 0.7067 |

For groups that differed based on developmental experience, we found pronounced coexpression differences in both drainages but in generally opposite directions: in the Aripo drainage more complex subnetwork motifs were more abundant in fish reared without predator cues while in the Quare drainage complex motifs were more abundant in fish reared with predators (Figure 4, Table 3). These patterns were largely concordant for DE and NDE gene sets within both drainages.

Population divergence in coexpression networks was also apparent, again with distinct patterns in the two lineages. In the Aripo drainage, high-predation fish from both rearing environments had generally fewer complex network motifs among NDE genes and no differences among DE genes (Figure 4, Table 3). In the Quare drainage, this pattern was flipped, with more complex motifs among NDE genes in high-predation fish, and more complex motifs among DE genes in low-predation fish (Figure 4, Table 3).

Differences between the two drainages were also pronounced, with the Aripo drainage having overall fewer complex network motifs, apart from a few differences in the opposing direction for triangles and 3-star motifs among NDE genes (Figure 4, Table 4).

## DISCUSSION

Our goal in this study was to understand how genetic background and rearing environment shape relationships among genes. We previously characterized expression changes at the level of individual genes (Fischer *et al*., 2021), and here we were interested in exploring changes in coexpression patterns among genes. Our findings suggest that coexpression patterns are flexible at evolutionary and developmental timescales. Exciting from a biological perspective, these questions present statistical challenges. We discuss the implications of our work from both angles.

### Overcoming sample size constraints in high-dimensional data

Gene expression studies remain plagued by small per-group samples sizes and high dimensionality. Network construction is far from trivial, if not problematic, under these conditions, especially when network structure – and not just network expression level – differs among experimental groups. In our own study, we had an overall sample size of N=98 individuals, well above the recommendation of N=30 for network construction. However, this total sample size includes samples from two drainages and four experimental groups, and – based on our analyses here and preliminary analyses using the WGCNA package – we found evidence that network structure differs between experimental groups and even more strongly between drainages. These differences are of key biological interest as they suggest that expression relationships among genes (i.e., network structure) are subject to developmental plasticity and evolutionary divergence. However, if network structure differs across experimental groups, then networks must be constructed separately for each experimental group to avoid construction of ‘average’ networks that can obscure differences of (biological) interest and lead to biased conclusions (Zhao *et al*., 2014; Shojaie, 2021; Li *et al*., 2022; Sai Li & Li, 2023). To take an extreme example, if two genes have opposing correlations of the same magnitude in two groups, the average correlation across groups will be zero. Thus, it is the per-group sample size that is most important for network construction and comparison when gene coexpression patterns are of interest.

While our per-group sample size of N=10-15 is relatively large for an RNAseq study, it is below the recommended threshold for network construction, such as the minimum sample of N=20 suggested for RNAseq analyses by (Langfelder & Horvath, 2008; Ballouz *et al*., 2015). As we illustrate, these sample sizes are surprisingly inadequate for recently developed statistical tests thought to be robust against high-dimensionality, to control Type I error, and to maintain power. Indeed, from our simulation experiments, most common methods require N>50 to retain the generally accepted nominal significance levels of 0.05 and satisfactory power exceeding 0.8. Importantly, the potential misinterpretations resulting from these shortcomings are not systematic (i.e., directionally biased) and therefore difficult to predict.

To address the challenges in two-sample covariance testing, we employed a random projection-based test rather than existing methods grounded in large-sample asymptotic theory. As shown in Supplementary Information B, our method effectively controls the type-I error rate and maintains reasonable power across various choices of projection dimensions and numbers of random projections, even when the sample size is as small as N=10. The test results reveal statistically significant differences in the covariance matrices for several group pairs, indicating structural differences in their underlying networks. Our statistical approach uses non–data-driven projections, which offer several advantages: (i) unlike data-driven methods such as PCA or random-skewers or eigentensor decomposition-biased approaches (Wang *et al*. 2019; Hu *et al*. 2025), the random projection-based test naturally preserves the null hypothesis; (ii) in the resulting low-dimensional projected space, powerful and robust test statistics, such as the U-statistics considered in this paper, can be easily constructed; (iii) multiple random projections yield conditionally independent tests, whose aggregation (e.g., via the maximum operator in this work) enhances power. While we demonstrate the utility of this procedure through simulations, a formal theoretical investigation of these advantages is warranted and is left for future statistical research. As a growing number of studies consider how interactions among genes shape phenotypic differences across timescales, we present our work as a case study to increase awareness of these limitations, present complementary statistical approaches to those commonly used, and in hopes that others will consider these issues in experimental design and analysis.

### Evidence for genetic and developmental differences in gene coexpression

Using the robust estimation methods we derived, we first identified differences based on both genetic background and developmental environment, with the most pronounced differences in both drainages between high- and low-predation fish reared with predators (HP pred+ vs LP pred+). This comparison represents the ancestral population adapted to life with predators (HP pred+) versus the derived low-predation population adapted to predator-free environments and suddenly re-exposed to predator cues (e.g., as when fish are washed downstream; LP pred+). Fish adapted to a low-predation life are poorly equipped deal with the sudden stressors of predation. Indeed, we previously found HP pred+ fish to be behaviorally least variable and LP pred+ fish to be behaviorally most variable – both in single behaviors and in the correlations among them (Fischer *et al*., 2016b). In light of findings here, we suggest that disruption of gene coexpression networks could contribute to unpredictable behavioral patterns and correlations.

Beyond genetic and developmental differences within each evolutionary lineage, we found that differences were also ubiquitous when comparing between the two lineages, and more evidence for coexpression differences in the Quare as compared to the Aripo drainage. We suggest these patterns arise in part from the extent of genetic divergence between populations: high- and low-predation populations in the Quare drainage show greater genetic (Willing *et al*., 2010a) and gene expression (Fischer *et al*., 2021) divergence than those in the Aripo drainage, and the two drainages represent distinct evolutionary lineages (Willing *et al*., 2010a). The importance of genetic background in shaping evolutionary trajectories is highlighted by our previous work demonstrating distinct underlying mechanisms associated with parallel phenotypic adaptation in guppies from distinct evolutionary lineages (Fischer *et al*., 2021). Similar mechanistic flexibility has also been demonstrated in other systems (Cordero *et al*., 2018; Jacobs *et al*., 2020), including those known for parallel phenotypic evolution (e.g. (Laporte *et al*., 2015; Hanson *et al*., 2017; Bolnick *et al*., 2018)). Our findings here extend these observations from the expression of individual genes to coexpression patterns among genes, suggesting that alternative gene expression network configurations can give rise to shared organism-level phenotypes.

The prevalence of coexpression differences in our datasets highlights the need to construct coexpression networks individually across experimental groups. If connectivity diverges at evolutionary timescales and shifts with developmental experience, then constructing a single network from pooled samples may confound differential expression differences with differences in connectivity. For example, if a subset of DE genes has higher connectivity in experimental group A versus experimental group B, then connectivity patterns associated with differential gene expression may be obscured when an average network with intermediate connectivity is constructed. Conversely, and non-mutually exclusive, genes differentially expressed between groups A and B could lead to the appearance of strong expression correlation (i.e., strong connectivity) overall, when in fact the within treatment correlations are weak. In brief, pooling samples for network construction when underlying networks in fact differ across experimental groups may obscure precisely the differences researchers are interested in testing.

We note that divergence in gene expression and by extension co-expression reflects both adaptive processes as well as non-adaptive and neutral processes (Whitehead & Crawford, 2006; Lynch, 2007). First, low-predation populations in guppies are particularly susceptible to drift, founder effects, and inbreeding depression because they are established by a small number of individuals and experience relaxed predator selection (Magurran, 2005b; Barson *et al*., 2009; Willing *et al*., 2010b). Second, whether initial gene expression changes are dominated by adaptive or non-adaptive processes, further expression changes may be homeostatic or compensatory, buffering higher-level phenotypes from evolutionary change, and thereby leading to alternative transcriptional configurations associated with shared organismal phenotypes (e.g., (Abouheif & Wray, 2002; Crawford & Oleksiak, 2007; Fischer *et al*., 2016b)). Finally, some changes in coexpression may simply reflect transcriptional noise that is filtered out by downstream regulatory processes (e.g., translational regulation). Though certainly present in our dataset, transcriptional noise is an unlikely a major driver of our results as random expression changes would need to be concordant across individuals to be detected as changes in coexpression. At present, we cannot distinguish between adaptive, non-adaptive, homeostatic, and neutral processes. Indeed, a combination of all factors shapes gene expression and coexpression, and distinguishing among alternatives remains a challenge in transcriptomic work. Developing high-dimensional methods such as ours to detect coexpression differences in small sample sizes is an important first step in understanding the causes and consequences of coexpression changes across developmental and evolutionary timescales.

### Probing links between network coexpression differences, developmental plasticity, and evolutionary divergence

By comparing the prevalence subgraph motifs across networks, we were able to make comparisons of network connectivity and clusterability among experimental groups to further characterize the coexpression differences. Our results indicate widespread differences in gene coexpression network structure (both connectivity and clusterability) at developmental and evolutionary timescales; however, these differences do not follow a directional rule. Prior work proposed alternative directional hypotheses, (1) genes in central network positions are evolutionarily constrained and hence less likely to diverge in mean expression levels; and (2) expression changes in genes occupying central hub positions are more likely to have phenotypic consequences and hence more likely to be targets of selection. Rather than supporting either consistently increased or decreased connectivity or clusterability, our findings provide a potential explanation for conflicting evidence surrounding the ongoing debate about whether hub genes are more or less likely to evolve: if coexpression relationships themselves can change both on developmental and short-term evolutionary time scales, then the constraints and/or advantages imposed by high versus low connectivity are not fixed. As a result, the association between network position and differential expression may vary across traits, taxa, and environments and therefore studies.

We detected extensive plasticity and divergence in connectivity and clusterability in both DE and NDE gene sets, highlighting that network changes do not have a simple association with differential expression levels of individual genes. We analyzed DE and NDE networks separately with the idea that divergence and plasticity in mean expression levels of DE genes might be driving the covariance differences we report. As a simplest example, covariance between two genes in one experimental group might be disrupted by the molecular or cellular mechanisms that increase or decrease the expression level of one of those genes during population divergence. Our results in Tables 3 and 4 refute the notion that differential gene expression is the primary driver of network differences, as even the genes that lacked detectable differences in mean expression levels (NDE genes) exhibited widespread divergence in coexpression subgraphs. This pattern raises key questions about the sources of coexpression flexibility, prompting future work analyzing the molecular and cellular mechanisms that drive these overall patterns, including changes in transcriptional regulation, mRNA stability, and changing proportions of cell types.

## Conclusions

Understanding how underlying genetic architecture shapes the maintenance and evolution of complex traits is a fundamental goal of biological research. Over the past two decades, the explosion of next-generation sequencing technologies has allowed us to move beyond the genetic scale – considering one or a few genes or loci – to genomic scales – considering thousands to tens of thousands of genes or loci. Among the key advances afforded by these approaches are the ease of conducting broadscale, exploratory studies; the opportunity to characterize underlying mechanisms in non-model species; and the ability to consider genes in the context of their interactions. As a growing number of studies consider how interactions among genes shape phenotypic differences across timescales, we provide a case study to increase awareness of limitations and provide suggestions for analysis.

Our findings provide intriguing evidence of extensive coexpression differences at multiple timescales in a species known for rapid adaptation and suggest that flexibility in gene coexpression relationships across time scales may contribute to evolutionary potential. This idea, its generality, and its consequences for adaptation will be revealed by more studies with larger sample sizes and new statistical approaches. We further identify divergence and plasticity in co-expression of genes that do not show differential expression levels, a pattern that raises many questions about the mechanistic basis of coexpression patterns and their divergence. We argue that pooling samples across experimental groups may obscure precisely the differential expression and connectivity differences that we seek to understand. Understanding whether and how relationships among genes change at developmental and evolutionary timescales has consequences for our understanding of how underlying mechanisms shape flexibility and robustness in higher order phenotypes, how animals adapt to novel and changing environments, and how behavior and physiology are regulated in health and disease.

## ACKNOWLEDGEMENTS

We thank the members of the Colorado State University Guppy Group for fish care and help with tissue collection and processing. We thank two reviewers for thoughtful comments that greatly improved the manuscript. This work was supported by the National Science Foundation DDIG-1311680 (to EKF), RCN IOS-1256839 (to EKF), IOS-1354755 (to KLH), IOS 1922701 (to WZ) , US Department of Energy DE-SC0018344 (to WZ), and NIH R01GM144961 (to WZ).

## DATA ACCESSIBILITY

Raw sequencing reads are available through the NCBI SRA repository (PRJNA601479). R code for statistical analyses will be made available on GitHub (https://github.com/EnigmaSong/GeneFlexibilityStudy) upon publication.

## AUTHOR CONTRIBUTIONS

EKF and KLH conceived of the study; EKF collected samples and performed molecular work, gene expression mapping, transcript abundance estimation, and preliminary differential expression analyses; YS and WZ devised and performed statistical analyses with input from EKF and KLH; EKF and YS performed data visualization; EKF wrote the manuscript with contributions from all authors.

## Notes

https://github.com/EnigmaSong/GeneFlexibilityStudy

https://www.ncbi.nlm.nih.gov/sra/

### Competing Interest Statement

The authors have declared no competing interest.

### Summary of Updates

We have made extensive revisions to the manuscript based on reviewer feedback. This includes additional analyses and figures, and changes to the text.

## REFERENCES

Abouheif, E. & Wray, G. 2002. Evolution of the gene network underlying wing polymorphism in ants. Science (1979) 297: 249–252.

Agterberg, J., Tang, M. & Priebe, C.E. 2020. Nonparametric two-sample hypothesis testing for random graphs with negative and repeated eigenvalues. arXiv 2012.09828.

Alyakin, A.A., Agterberg, J., Helm, H.S. & Priebe, C.E. 2024. Correcting a Nonparametric Two-sample Graph Hypothesis Test for Graphs with Different Numbers of Vertices with Applications to Connectomics. Appl Netw Sci 9: 1.

Badyaev, A. V. 2018. Evoulutionary transitions in controls reconcile adaptation with continuity of evolution. Semin Cell Dev Biol 88: 36–45.

Ballouz, S., Verleyen, W. & Gillis, J. 2015. Guidance for RNA-seq co-expression network construction and analysis: Safety in numbers. Bioinformatics 31.

Barson, N.J., Cable, J. & Van Oosterhout, C. 2009. Population genetic analysis of microsatellite variation of guppies (*Poecilia reticulata*) in Trinidad and Tobago: Evidence for a dynamic source-sink metapopulation structure, founder events and population bottlenecks. J Evol Biol 22: 485–497.

Bolnick, D.I., Barrett, R.D.H., Oke, K.B., Rennison, D.J. & Stuart, Y.E. 2018. (Non)Parallel evolution. Annu Rev Ecol Evol Syst 49: 303–330.

Cai, T., Liu, W. & Xia, Y. 2013. Two-sample covariance matrix testing and support recovery in high-dimensional and sparse settings. J Am Stat Assoc 108: 265–277.

Cai, T.T. & Liu, W. 2016. Large-Scale Multiple Testing of Correlations. J Am Stat Assoc 111: 229– 240.

Chang, J., Zhou, W., Zhou, W.X. & Wang, L. 2017. Comparing large covariance matrices under weak conditions on the dependence structure and its application to gene clustering. Biometrics 73: 31–41.

Chateigner, A., Lesage-Descauses, M.C., Rogier, O., Jorge, V., Leplé, J.C., Brunaud, V., et al. 2020. Gene expression predictions and networks in natural populations supports the omnigenic theory. BMC Genomics 21: 1–16. BMC Genomics.

Chowdhury, H.A., Bhattacharyya, D.K. & Kalita, J.K. 2020. (Differential) Co-Expression Analysis of Gene Expression: A Survey of Best Practices. IEEE/ACM Trans Comput Biol Bioinform 17: 1154–1173. IEEE.

Cordero, G.A., Liu, H., Wimalanathan, K., Weber, R., Quinteros, K. & Janzen, F.J. 2018. Gene network variation and alternative paths to convergent evolution in turtles. Evol Dev 20: 172– 185.

Crawford, D.L. & Oleksiak, M.F. 2007. The biological importance of measuring individual variation. Journal of Experimental Biology 210: 1613–1621.

Des Marais, D.L., Guerrero, R.F., Lasky, J.R. & Scarpino, S. V. 2017. Topological features of a gene co-expression network predict patterns of natural diversity in environmental response. Proceedings of the Royal Society B: Biological Sciences 284: 20170914.

Endler, J.A. 1995. Multiple-trait coevolution and environmental gradients in guppies. Trends Ecol Evol 10: 22–29.

Fischer, E.K., Ghalambor, C.K. & Hoke, K.L. 2016a. Can a Network Approach Resolve How Adaptive vs Nonadaptive Plasticity Impacts Evolutionary Trajectories? Integr Comp Biol 56: 877–888.

Fischer, E.K., Ghalambor, C.K. & Hoke, K.L. 2016b. Plasticity and evolution in correlated suites of traits. J Evol Biol 29: 991–1002.

Fischer, E.K., Harris, R.M., Hofmann, H.A. & Hoke, K.L. 2014. Predator exposure alters stress physiology in guppies across timescales. Horm Behav 65: 165–172. Elsevier Inc.

Fischer, E.K., Soares, D., Archer, K.R., Ghalambor, C.K. & Hoke, K.L. 2013. Genetically and environmentally mediated divergence in lateral line morphology in the Trinidadian guppy (*Poecilia reticulata*). Journal of Experimental Biology 216: 3132–3142.

Fischer, E.K., Song, Y., Hughes, K.A., Zhou, W. & Hoke, K.L. 2021. Non-parallel transcriptional divergence during parallel adaptation. Mol Ecol 30: 1516–1530.

Fraser, B.A., Künstner, A., Reznick, D.N., Dreyer, C. & Weigel, D. 2015. Population genomics of natural and experimental populations of guppies (*Poecilia reticulata*). Mol Ecol 24: 389–408.

Friedman, D.A., York, R.A., Hilliard, A.T. & Gordon, D.M. 2020. Gene expression variation in the brains of harvester ant foragers is associated with collective behavior. Commun Biol 3. Springer US.

Ghalambor, C.K., Hoke, K.L., Ruell, E.W., Fischer, E.K., Reznick, D.N. & Hughes, K.A. 2015. Non-adaptive plasticity potentiates rapid adaptive evolution of gene expression in nature. Nature 525: 372–375.

Ghalambor, C.K., McKay, J.K., Carroll, S.P. & Reznick, D.N. 2007. Adaptive versus non-adaptive phenotypic plasticity and the potential for contemporary adaptation in new environments. Funct Ecol 21: 394–407.

Gilliam, J.F., Fraser, D.F. & Alkins-Koo, M. 1993. Structure of a Tropical Stream Fish Community : A Role for Biotic Interactions. Ecology 74: 1856–1870.

Hahn, M.W., Conant, G.C. & Wagner, A. 2004. Molecular Evolution in Large Genetic Networks: Does Connectivity Equal Constraint? J Mol Evol 58: 203–211.

Hahn, M.W. & Kern, A.D. 2005. Comparative genomics of centrality and essentiality in three eukaryotic protein-interaction networks. Mol Biol Evol 22: 803–806.

Hämälä, T., Gorton, A.J., Moeller, D.A. & Tiffin, P. 2020. Pleiotropy facilitates local adaptation to distant optima in common ragweed (*Ambrosia artemisiifolia*). PLoS Genet 16: 1–23.

Handelsman, C.A., Broder, E.D., Dalton, C.M., Ruell, E.W., Myrick, C.A., Reznick, D.N., et al. 2013. Predator-induced phenotypic plasticity in metabolism and rate of growth: Rapid adaptation to a novel environment. Integr Comp Biol 53: 975–988.

Handelsman, C.A., Ruell, E.W., Torres-Dowdall, J. & Ghalambor, C.K. 2014. Phenotypic Plasticity Changes Correlations of Traits Following Experimental Introductions of Trinidadian Guppies (*Poecilia reticulata*). Integr Comp Biol 54: 794–804.

Hanson, D., Hu, J., Hendry, A.P. & Barrett, R.D.H. 2017. Heritable gene expression differences between lake and stream stickleback include both parallel and antiparallel components. Heredity (Edinb) 119: 339–348.

Harnqvist, S. 2021. Network centrality is the most important variable determining gene sequence evolution rates in *Schizosaccharomyces pombe*. bioRxiv 2021.01.10.

Hoke, K.L., Adkins-Regan, E., Bass, A.H., McCune, A.R. & Wolfner, M.F. 2019. Co-opting evo-devo concepts for new insights into mechanisms of behavioural diversity. Journal of Experimental Biology 222: jeb190058.

Hu, J., Weber, J.N., Fuess, L.E., Steinel, N.C., Bolnick, D.I. and Wang, M., 2025. A spectral framework to map QTLs affecting joint differential networks of gene co-expression. PLOS Computational Biology, 21(4): e1012953.

Huizinga, M., Ghalambor, C.K. & Reznick, D.N. 2009. The genetic and environmental basis of adaptive differences in shoaling behaviour among populations of Trinidadian guppies, *Poecilia reticulata*. J Evol Biol 22: 1860–1866.

Jacobs, A., Carruthers, M., Yurchenko, A., Gordeeva, N. V., Alekseyev, S.S., Hooker, O., et al. 2020. Parallelism in eco-morphology and gene expression despite variable evolutionary and genomic backgrounds in a Holarctic fish. PLOS Genetics. 16(4): e1008658.

Jeong, H., Mason, S.P., Barabási, A.L. & Oltvai, Z.N. 2001. Lethality and centrality in protein networks. Nature 411: 41–42.

Jin, X., Chan, K., Barnett, I. & Ghosh, R.P. 2024. Two-Sample Hypothesis Testing for Large Random Graphs of Unequal Size. *arXiv* 2402:11133.

Josephs, E.B., Wright, S.I., Stinchcombe, J.R. & Schoen, D.J. 2017. The relationship between selection, network connectivity, and regulatory variation within a population of *Capsella grandiflora*. Genome Biol Evol 9: 1099–1109.

Jovelin, R. & Phillips, P.C. 2009. Evolutionary rates and centrality in the yeast gene regulatory network. Genome Biol 10: R35.

Kim, P.M., Korbel, J.O. & Gerstein, M.B. 2007. Positive selection at the protein network periphery: Evaluation in terms of structural constraints and cellular context. Proc Natl Acad Sci U S A 104: 20274–20279.

Koubkova-Yu, T.C.T., Chao, J.C. & Leu, J.Y. 2018. Heterologous Hsp90 promotes phenotypic diversity through network evolution. PLoS Biol 16(11): e2006450.

Kuo, H.C., Yao, C. Te, Liao, B.Y., Weng, M.P., Dong, F., Hsu, Y.C., et al. 2023. Weak gene–gene interaction facilitates the evolution of gene expression plasticity. BMC Biol 21: 1–20.

Langfelder, P. & Horvath, S. 2008. WGCNA: An R package for weighted correlation network analysis. BMC Bioinformatics 9: 1–13.

Langfelder, P., Luo, R., Oldham, M.C. & Horvath, S. 2011. Is my network module preserved and reproducible? PLoS Comput Biol 7(1): e1001057.

Laporte, M., Rogers, S.M., Dion-Côté, A.M., Normandeau, E., Gagnaire, P.A., Dalziel, A.C., et al. 2015. RAD-QTL mapping reveals both genome-level parallelism and different genetic architecture underlying the evolution of body shape in lake Whitefish (*Coregonus clupeaformis*) species pairs. G3: Genes, Genomes, Genetics 5: 1481–1491.

Li, J. & Chen, S.X. 2012. Two sample tests for high-dimensional covariance matrices. Ann Stat 40: 908–940.

Li, S., Cai, T.T. & Li, H. 2022. Transfer Learning in Large-Scale Gaussian Graphical Models with False Discovery Rate Control. J Am Stat Assoc 118(543): 2171–2183.

Love, M.I., Huber, W. & Anders, S. 2014. Moderated estimation of fold change and dispersion for RNA-seq data with DESeq2. Genome Biol 15: 1–21.

Luisi, P., Alvarez-Ponce, D., Pybus, M., Fares, M.A., Bertranpetit, J. & Laayouni, H. 2015. Recent positive selection has acted on genes encoding proteins with more interactions within the whole human interactome. Genome Biol Evol 7: 1141–1154.

Lynch, M. 2007. The evolution of genetic networks by non-adaptive processes. Nat Rev Genet 8: 803–813.

Magurran, A.E. 2005a. Evolutionary ecology: the Trinidadian guppy. Oxford University Press, Oxford.

Mähler, N., Wang, J., Terebieniec, B.K., Ingvarsson, P.K., Street, N.R. & Hvidsten, T.R. 2017. Gene co-expression network connectivity is an important determinant of selective constraint. PLoS Genet 13: 1–33.

Masalia, R.R., Bewick, A.J. & Burke, J.M. 2017. Connectivity in gene coexpression networks negatively correlates with rates of molecular evolution in flowering plants. PLoS One 12: 1– 10.

Maugis, P.-A.G., Olhede, S.C. & Wolfe, P.J. 2017. Topology Reveals Universal Features for Network Comparison. *arXiv* 1705.05677.

McGuigan, K. & Sgrò, C.M. 2009. Evolutionary consequences of cryptic genetic variation. Trends Ecol Evol 24: 305–311.

Paaby, A.B. & Rockman, M. V. 2014. Cryptic genetic variation: Evolution’s hidden substrate. Nat Rev Genet 15: 247–258.

Pavličev, M. & Cheverud, J.M. 2015. Constraints Evolve: Context Dependency of Gene Effects Allows Evolution of Pleiotropy. Annu Rev Ecol Evol Syst 46: 413–434.

Pavlicev, M. & Wagner, G.P. 2012. A model of developmental evolution: Selection, pleiotropy and compensation. Trends Ecol Evol 27: 316–322.

Qi, M., Li, T. & Zhou, W. 2024. Multivariate Inference of Network Moments by Subsampling. *arXiv* 2409.01599.

Qiu, T., Xu, W. & Zhu, L. 2021. Two-sample test in high dimensions through random selection. Comput Stat Data Anal 160: e107218.

Rennison, D.J. & Peichel, C.L. 2022. Pleiotropy facilitates parallel adaptation in sticklebacks. Mol Ecol 31: 1476–1486.

Reznick, D., Butler IV, M.J. & Rodd, H. 2001. Life-History Evolution in Guppies. VII. The Comparative Ecology of High-and Low-Predation Environments. Am Nat 157: 126–140.

Reznick, D.A., Bryga, H. & Endler, J.A. 1990. Experimentally induced life-history evolution in a natural population. Nature 346: 357–359.

Reznick, D.N. 1997. Life history evolution in guppies (*Poecilia reticulata*): Guppies as a model for studying the evolutionary biology of aging. Exp Gerontol 32: 245–258.

Ruell, E.W., Handelsman, C.A., Hawkins, C.L., Sofaer, H.R., Ghalambor, C.K. & Angeloni, L. 2013. Fear, food and sexual ornamentation: plasticity of colour development in Trinidadian guppies. Proceedings of the Royal Society B: Biological Sciences 280: 20122019– 20122019.

Sai Li, T.T.C. & Li, H. 2023. Transfer Learning in Large-Scale Gaussian Graphical Models with False Discovery Rate Control. J Am Stat Assoc 118: 2171–2183.

Shao, M., Xia, D., Zhang, Y., Wu, Q. & Chen, S. 2022. Higher-Order Accurate Two-Sample Network Inference and Network Hashing. arXiv 2208.07573.

Shojaie, A. 2021. Differential network analysis: A statistical perspective. WIRES Compt Stat 13: e1508.

Storey, J.D. 2002. A direct approach to false discovery rates. J R Stat Soc Series B Stat Methodol 64: 479–498.

Strimmer, K. 2008. fdrtool: A versatile R package for estimating local and tail area-based false discovery rates. Bioinformatics 24: 1461–1462.

Tang, M., Athreya, A., Sussman, D.L., Lyzinski, V., Park, Y. & Priebe, C.E. 2017. A Semiparametric Two-Sample Hypothesis Testing Problem for Random Graphs. Journal of Computational and Graphical Statistics 26: 344–54.

Tommasini, D. & Fogel, B.L. 2023. multiWGCNA: an R package for deep mining gene co-expression networks in multi-trait expression data. BMC Bioinformatics 24: 1–15. BioMed Central.

Torres Dowdall, J., Handelsman, C.A., Ruell, E.W., Auer, S.K., Reznick, D.N. & Ghalambor, C.K. 2012. Fine-scale local adaptation in life histories along a continuous environmental gradient in Trinidadian guppies. Funct Ecol 26: 616–627.

Torres-Dowdal, J., Handelsman, C. a, Reznick, D.N. & Ghalambor, C.K. 2012. Local adaptation and the evolution of of phenotypic plasticity in Trinidadian guppies (*Poecilia reticulata*). Evolution 66: 3432–3443.

Wang, D., Wang, J., Jiang, Y., Liang, Y. & Xu, D. 2017. BFDCA: A Comprehensive Tool of Using Bayes Factor for Differential Co-Expression Analysis. J Mol Biol 429: 446–453. Elsevier Ltd.

Wang, Z., Liao, B.-Y. & Zhang, J. 2010. Genomic patterns of pleiotropy and the evolution of complexity. Proc Natl Acad Sci 107: 18034–18039.

Wang, M., Fischer, J. and Song, Y.S. 2019. Three-way clustering of multi-tissue multi-individual gene expression data using semi-nonnegative tensor decomposition. Annals of Applied Statistics, 13(2): 1103–1127.

Warnefors, M. & Kaessmann, H. 2013. Evolution of the correlation between expression divergence and protein divergence in mammals. Genome Biol Evol 5: 1324–1335.

West-Eberhard, M.J. 2003. Developmental plasticity and evolution. Oxford University Press, New York.

Whitehead, A. & Crawford, D.L. 2006. Neutral and adaptive variation in gene expression. Proc Natl Acad Sci 103(14): 5425–5430.

Willing, E.M., Bentzen, P., Van Oosterhout, C., Hoffmann, M., Cable, J., Breden, F., et al. 2010a. Genome-wide single nucleotide polymorphisms reveal population history and adaptive divergence in wild guppies. Mol Ecol 19: 968–984.

Willing, E.M., Bentzen, P., Van Oosterhout, C., Hoffmann, M., Cable, J., Breden, F., et al. 2010b. Genome-wide single nucleotide polymorphisms reveal population history and adaptive divergence in wild guppies. Mol Ecol 19: 968–984.

Wu, T.-L. & Li, P. 2020. Projected Tests for High-Dimensional Covariance Matrices. J Stat Plan Inference 207: 73–85.

Yu, X., Li, D. & Xue, L. 2020. Fisher’s Combined Probability Test for High-Dimensional Covariance Matrices. arXiv 2006.00426.

Zhao, S.D., Cai, T.T. & Li, H. 2014. Direct estimation of differential networks. Biometrika 101: 253–268.

