## Supplemental Materials for "Flexibility in Gene Coexpression at Developmental and Evolutionary Timescales"

Eva K. Fischer

Youngseok Song

Wen Zhou

Kim L. Hoke

### SUMMARY OF PRELIMINARY ANALYSES USING WGCNA

*Methods*

Our initial approach was to characterize and explore (differences in) gene coexpression using the popular package Weighted Gene Correlation Network Analysis (WGCNA) package in R (Langfelder & Horvath, 2008). WGCNA constructs gene modules based on coexpression patterns. Module construction is unsupervised and does not require previous knowledge of gene-gene relationships; however, sufficient sample size (N=20) is critical to obtain the approximately scale-free network topology on which module construction relies. We reasoned that this approach was therefore only feasible if networks were preserved across groups, such that all samples could be used for construction of a single set of shared modules. This was particularly important given our interest in whether the relationships among genes, rather than solely the expression of individuals genes, was altered by developmental and evolutionary influences.

To make approaches most comparable, we included all genes that passed our filtering criteria in WGCNA analysis (Aripo: 13,446 ; Quare: 14,379). Our filtering criteria are in line with the package recommendations to restrict analyses to the most variable genes, although we are more liberal in our variability threshold and therefore include a relatively large number of genes in network construction.

We constructed networks following general recommendations and the WGCNA tutorial for working with multiple datasets. We used a soft-thresholding power of 10 and a minimum module size of 30. Networks were constructed and modules identified separately for each experimental group (HP pred-, HP pred+, LP pred-, LP pred+) within each drainage. Following initial network construction, we merged modules based on similarities in module expression, as per tutorial recommendations. We report results for final, merged modules. We compared module similarity between experimental groups within drainage using the module preservation approach provided by the developers (Langfelder, Luo, Oldham, & Horvath, 2011). To compare module preservation across groups of interest, WGCNA calculates a composite $Z_{summary}$ value. $Z_{summary}$ values >10 indicate high module preservation and >2 moderate module preservation. We calculated $Z_{summary}$ metrics and visualized module preservation by superimposing the clustering pattern for the HP pred- group onto all other groups while preserving the module assignments (i.e., colors) from their own experimental group.

*Results*

Using WGCNA, we identified modules for each experimental group in both drainages and found overall only moderate preservation across groups (Figure SA1). Comparing across all experimental groups, average module preservation was lower in the Aripo dataset (median $Z_{summary}$ = 5.77; 18/119 modules $Z_{summary}$ >10) than in the Quare dataset (median $Z_{summary}$ = 13.22; 11/19 modules $Z_{summary}$ >10). We additionally calculated a $Z_{summary}$ value for a module of N=500 randomly selected genes (the gold module) to provide an estimate of the degree of module preservation expected by chance. For the Aripo drainage, 70/119 (59%) modules had higher $Z_{summary}$ values than the gold module, and for the Quare drainage 8/19 (42%) modules had higher $Z_{summary}$ values than the gold module. Thus, although raw preservations scores were higher in Quare, the percentage of modules more preserved than expected by chance was not. Overall, we identified a larger number of on average smaller modules in the Aripo drainage than in the Quare drainage (Table SA1).

*Conclusions*

Taken together, WGCNA analyses indicate that – while some modules are preserved – approximately half of all modules differ among experimental groups, based on both preservation $Z_{summary}$ scores and comparisons to randomly constructed modules. These results point to the fact that there are substantial network differences between experimental groups. In other words, networks should be constructed separately for each group. While of biological interest, this recognition creates technical difficulties given small (N=10-12) per group sample sizes. This is issue we sought to address with alternative statistical approaches.

**Table SA1.** Module numbers and average sizes in each experimental group and drainage. Data are for modules after merging modules with similar expression profiles.

| | **Aripo drainage** | | **Quare drainage** | |
|---|---|---|---|---|
| | module # | median module size | module # | median module size |
| HP pred- | 118 | 71.5 [30 – 837] | 18 | 242.5 [58 – 3888] |
| HP pred+ | 118 | 73 [30 – 547] | 33 | 175 [46 – 2676] |
| LP pred - | 88 | 63.5 [32 – 1595] | 11 | 172 [30 – 7316] |
| LP pred+ | 126 | 68 [31 – 750] | 12 | 483 [57 – 4032] |

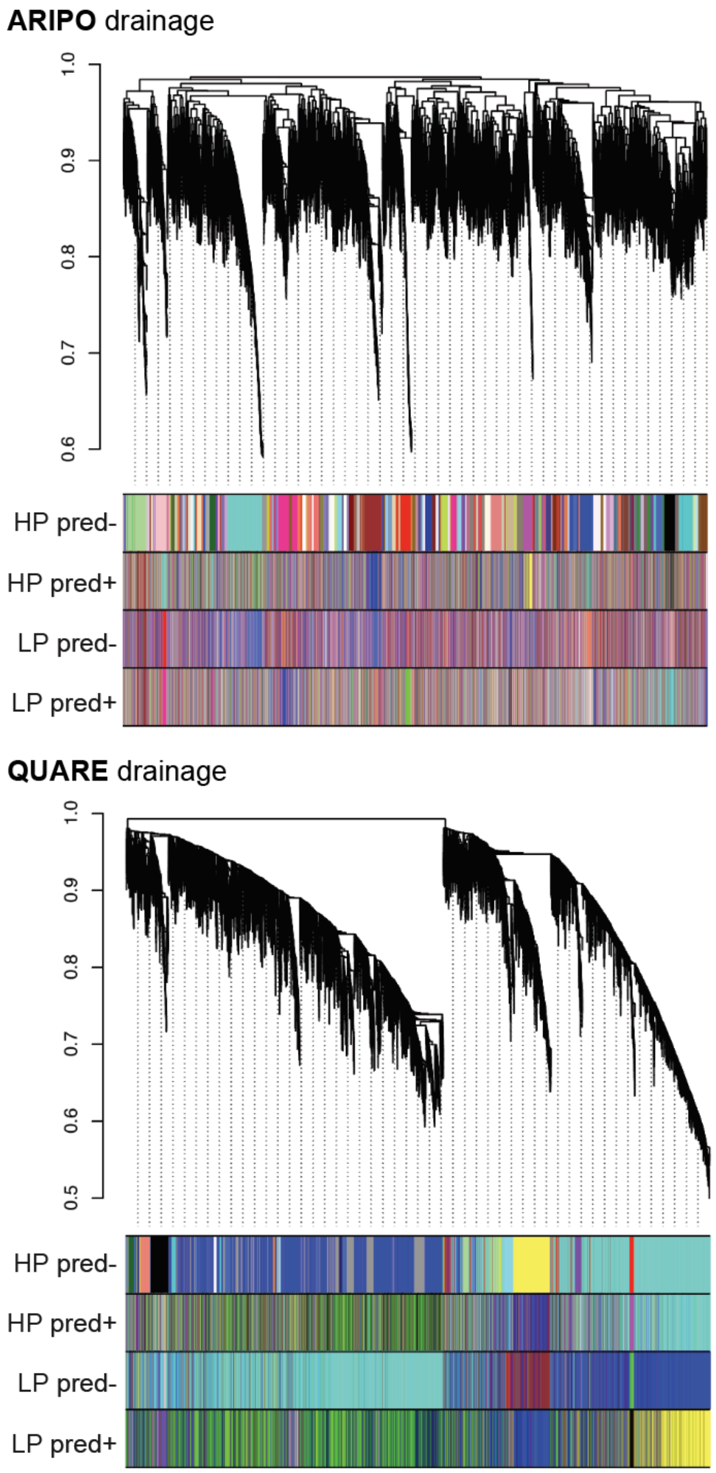


**Figure SA1.** Visualization of module (non-)preservation across groups. Gene clustering pattern for high-predation pred- (HP pred-; top row) is superimposed onto the other experimental groups while maintaining color-coded module assignments for each group. Scrambling of colors indicates that modules are not preserved. More similarity in color blocks across groups in the Quare drainage illustrates greater module preservation in this drainage. Note also the greater number of modules identified in the Aripo versus the Quare drainage (more colors overall in Aripo).
