## Supplemental Materials for "Flexibility in Gene Coexpression at Developmental and Evolutionary Timescales"

April 26, 2025

In the supporting information, we outline the statistical analysis procedures used in the paper, which include covariance matrix comparison methods and network comparison of two gene sets in the same group. First, addressing concerns from the small sample size in Section 2, we introduce an alternative method designed to enhance the power while controlling the type-I error rate in Section 3. Moreover, in Section 4.3, we compare gene networks with different numbers of nodes by network summary plots and a two-sample network test.

#### 1 Experimental Design and Data Preprocessing

We first describe the experimental design used in this manuscript, which was also used in Fischer et al. (2021). We used the $2 \times 2$ full-factorial two-way ANOVA as Table S1. Two river drainages represent parallel, independent evolutionary lineages (Aripo and Quare), each with two populations, high-predation (HP) and low-predation (LP), and two rearing environments, without chemical predator cures (pred-) and with chemical predator cues (pred+). Using these abbreviations, we label the groups within each drainage as HP pred+, HP pred-, LP pred+, and LP pred- for the rest of the supporting information. Table S2 displays the number of samples in each group. In the rest of the supporting document, we refer to the dataset from Aripo and Quare drainages as the *Aripo* and *Quare* datasets. The Aripo dataset is balanced, while the Quare dataset is not.

We ran data preprocessing for statistical analysis on covariance structure or correlation networks. We used the normalized counts from the differential gene expression analysis in

Table S1: Abbreviations for the group names.

| Population & Rearing condition | Predation (P) | Non-predation (NP) |
|---|---|---|
| Low-predation (LP) | LP pred+ | LP pred- |
| High-predation (HP) | HP pred+ | HP pred- |

Table S2: Number of samples of groups.

| | LP pred+ | LP pred- | HP pred+ | HP pred- | Total |
|---|---|---|---|---|---|
| Aripo | 10 | 10 | 10 | 10 | 40 |
| Quare | 15 | 16 | 15 | 12 | 58 |

Table S3: Number of genes used in covariance tests and network analysis. DE indicates differential expressed genes in the population effect obtained in Fischer et al. (2021).

| | DE | NDE | Total |
|---|---|---|---|
| Aripo | 553 | 12891 | 13447 |
| Quare | 2909 | 11470 | 14379 |

Fischer et al. (2021). From these datasets, we filtered genes with variances less than 100 as an ad hoc cut-off. Next, to remove the mean trend, we obtained residuals from the linear mixed effect model. Specifically, we used the log transformation of the gene expression data as the response variable, two fixed effects (population and rearing environment), and two random effects (family and week). In other words, we obtain the residual from

$$\begin{aligned} y_{ijk} &= \mu + \alpha_i + \beta_j + (\alpha\beta)_{ij} + a_{ijk} + b_{ijk} + \epsilon_{ijk}, \\ e_{ijk} &= y_{ijk} - \widehat{y}_{ijk}, \end{aligned} \tag{1.1}$$

where $y_{ijk}$ is the log-transformed normalized count, $\widehat{y}_{ijk}$ represents the predicted response from the model, $\alpha_i$ and $\beta_j$ represent the effect of population and the rearing, respectively, $(\alpha\beta)_{ij}$ is the interaction effect, and $a_{ijk}$ and $b_{ijk}$ are random intercepts from week and family, respectively. We used the preprocessed data aimed to compare covariance matrices and correlation networks. However, we encountered two challenges: one from a small sample size, and the other from unequal network sizes.

The first challenge arises from the small sample size and the large number of genes, a so-called high-dimensional scenario. Under this scenario, several methods have been proposed for two-sample covariance testing including Li and Chen (2012) and Cai et al. (2013). However, these standard methods use the asymptotic distribution of the test statistic, which means the methods rely on approximation from large-sample theory, and the type-I error rate of the tests is not controlled when the sample size is small. The sample sizes in our datasets for groups defined in Table S1 are in Table S2. These per-group sample sizes are smaller than 60, which is the smallest $n$ considered in the simulation studies from both Li and Chen (2012) and Cai et al. (2013).

The second challenge arises from the unequal sizes of the two correlation networks. We want to compare two correlation networks from differentially expressed (DE) genes and non-DE (NDE) genes in the population effect, obtained from Fischer et al. (2021). If two correlation networks have the same size, two-sample correlation tests can be applied. However, the sizes of the two networks in our case (DE and NDE) are different, as shown in Table S3. Thus standard two-sample correlation tests are not applicable. To overcome this issue, we employ network analysis methods to compare correlation networks with different numbers of nodes.

The rest of the supporting information is organized as follows. In Section 2, we demonstrate that standard high-dimensional two-sample covariance tests suffer from either inflated type-I error rate or lower power due to small sample size. Section 2.1 discusses about why applying the post-selection along do not resolve this type-I error rate issue. We discuss resolutions of this issue in Section 3 and introduce the random projection test with $\ell_2$-type test statistics and numerical simulation results. We provide analysis results using the real data in Section 4. We also provide extra numerical simulation results regarding the critical value generation in Appendix A.

**Notation.** The sample size and the dimension of each sample are denoted by $n$ and $p$, respectively. We denote matrices by bold capital letters such as $\mathbf{\Sigma}$ and $\mathbf{D}$. The matrix $\mathbf{I}$ represents the identity matrix. Vectors are denoted by bold lowercase letters, and $\mathbf{0}$ is a zero vector. Multivariate normal distribution with the mean vector $\boldsymbol{\mu}$ and the covariance matrix $\mathbf{\Sigma}$ is denoted by $N(\boldsymbol{\mu}, \mathbf{\Sigma})$. The $\ell_2$-norm of a vector $\mathbf{v} = (v_1, \ldots, v_p)^{\mathrm{T}}$ is defined by $\|\mathbf{v}\|_2 := \sqrt{\sum_{j=1}^{p} v_j^2}$.

### 2 Challenges from small sample size

To compare the connectivity patterns of two groups, we can use two-sample covariance tests. Given the scenario where the sample size $n$ is much smaller than the dimension of the data $p$, we should use a high-dimensional covariance test. However, if the sample size is smaller, as in our situation, tests would not control the type-I error rate and have reasonable power. In the rest of the section, we present empirical study results to illustrate this phenomenon.

We check the type-I error rate of the selected two-sample covariance tests: Li and Chen (2012) (LC), Cai et al. (2013) (CLX), and Yu et al. (2024) (FC) using simulated data. We consider varying sample sizes for two simulated sets, $n = n_1 = n_2 \in \{10, 15, 20, 30, 50, 100\}$, and the dimension of each sample vector, $p \in \{250, 500, 1000, 2500, 5000, 10000\}$. We generate $n_1 + n_2$ random samples from $N(\mathbf{0}, \mathbf{\Sigma})$ where $\mathbf{\Sigma} = \mathbf{D}\widetilde{\mathbf{\Sigma}}\mathbf{D}$, $\mathbf{D} = \mathrm{diag}(d_1, \ldots, d_p)$, $d_j \sim \mathrm{uniform}(1, 5)$, and $\widetilde{\mathbf{\Sigma}} = \{\widetilde{\sigma}_{j,k}\}_{1 \leqslant j,k \leqslant p}$ defined as follows:

1. (independent) $\widetilde{\mathbf{\Sigma}} = \mathbf{I}$.

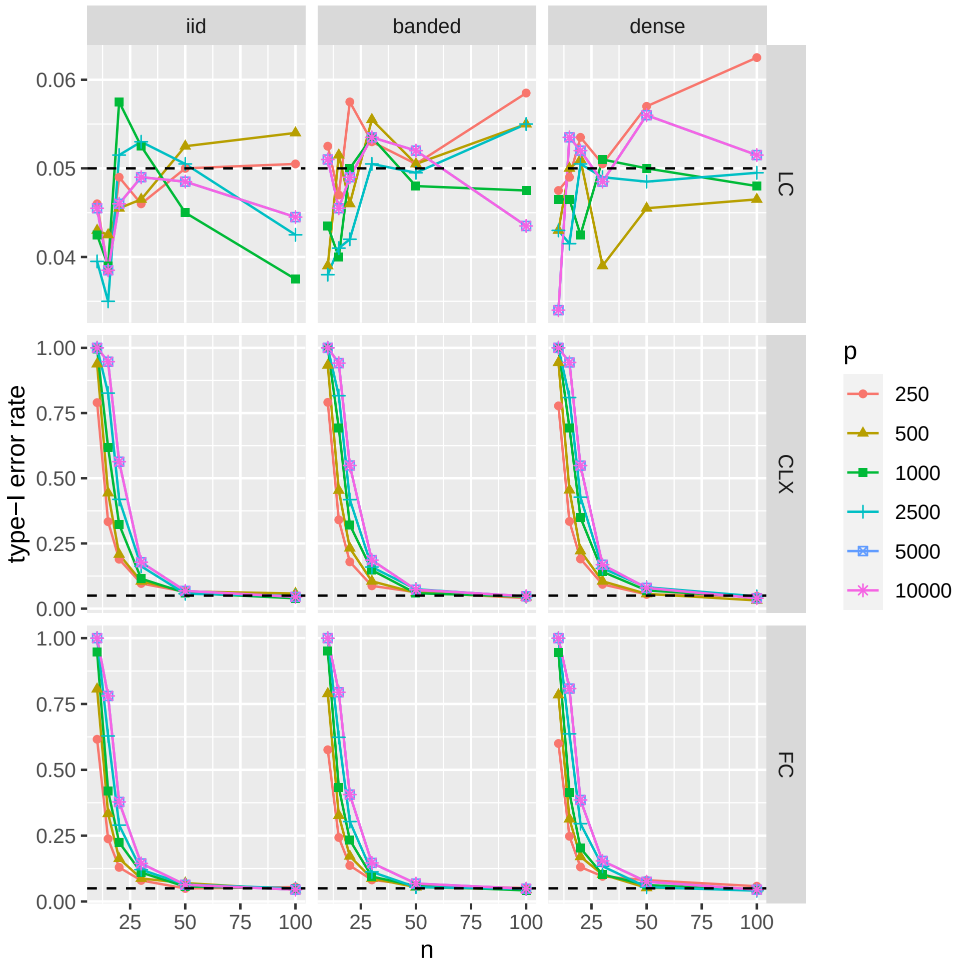


Figure S1: Type-I error rate of covariance tests, LC (Li and Chen, 2012), CLX (Cai et al., 2013), and FC (Yu et al., 2024) with the significance level of $\alpha = 0.05$, $p \in \{250, 500, 1000, 2500, 5000, 10000\}$ based on 2000 replications by the sample sizes $n = n_1 = n_2$.

2. (Banded) $\tilde{\sigma}_{j,j} = 1$, $\tilde{\sigma}_{j,j\pm1} = 0.6$, $\tilde{\sigma}_{j,j\pm2} = 0.3$, and $\tilde{\sigma}_{j,k} = 0$ when $|j - k| \geqslant 3$.

3. (Dense) $\tilde{\sigma}_{j,k} = 0.6^{|j-k|}$.

We use the asymptotic distribution of the test statistics for computing $p$-values and type-I error rate based on 2000 replications, using the significance level of $\alpha = 0.05$.

Figure S1 displays type-I error rates obtained from the simulation with a significance level of $\alpha = 0.05$. We observe that the LC test controls the type-I error rate for $n \leqslant 15$, while the CLX and FC tests do not, especially for the small sample size. This discrepancy arises from the limitation of asymptotic distribution approximation, which is based on large-sample theory, thus it may not work well for small sample size. In other words, the

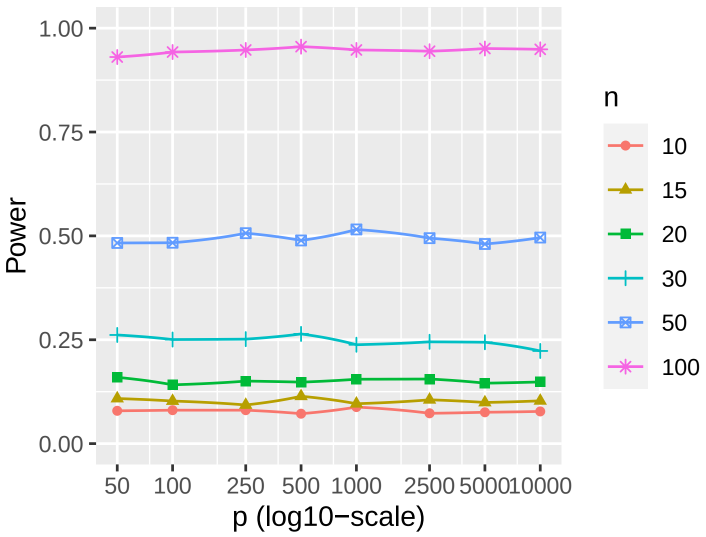


Figure S2: Empirical power of the LC test for $p \in \{50, 100, 250, 500, 1000, 2500, 5000, 10000\}$ and the covariance setting (2.1). The power is computed based on 2000 repetitions.

type-I error rate may not be controlled if the approximation has a large error. In summary, among the three tests considered, the LC test is the only one that controls the type-I error rate when $n \leqslant 15$, making it the primary candidate for consideration. However, we will later demonstrate that it lacks sufficient power under this scenario.

Next, we perform a power analysis of the LC test for a small sample size. Recall that controlling the type-I error rate does not guarantee that the test has sufficient power. To verify the power of the LC test for small sample sizes, we borrow one of the data generation settings from Wu and Li (2020). They considered two covariance matrices,

$$\mathbf{\Sigma}_1 = \begin{bmatrix} d_1 & \rho_1 & \rho_1^2 & \cdots & \rho_1^{p-1} \\ \rho_1 & d_1 & \rho_1 & \cdots & \rho_1^{p-2} \\ \rho_1^2 & \rho_1 & d_1 & \cdots & \rho_1^{p-3} \\ \vdots & \vdots & \vdots & \ddots & \vdots \\ \rho_1^{p-1} & \rho_1^{p-2} & \rho_1^{p-3} & \ddots & d_1 \end{bmatrix}, \mathbf{\Sigma}_2 = \begin{bmatrix} d_2 & \rho_2 & \rho_2^2 & \cdots & \rho_2^{p-1} \\ \rho_2 & d_2 & \rho_2 & \cdots & \rho_2^{p-2} \\ \rho_2^2 & \rho_2 & d_2 & \cdots & \rho_2^{p-3} \\ \vdots & \vdots & \vdots & \ddots & \vdots \\ \rho_2^{p-1} & \rho_2^{p-2} & \rho_2^{p-3} & \ddots & d_2 \end{bmatrix}, \tag{2.1}$$

where $(d_1, \rho_1, d_2, \rho_2) = (1.5, 0.5, 1, 0.6)$. The two covariance matrices in (2.1) are different overall, but the difference in each entry is small. Hence, we expected that the LC test would capture the difference, but it becomes more challenging when the sample size is small.

Figure S2 displays the empirical power of the LC test from 2000 repetitions. We observe the power of LC test is less than 0.25 when the sample size is smaller than 30. The dimension of the data matrix $p$ does not have much influence on the power of the test under these conditions. The message from Figure S2 is twofold. First, it raises concern over the power of the LC test for our dataset, which has a small sample size and large dimensionality

Table S4: Empirical type-I error rate of high-dimensional one-sample covariance tests with $H_0 : \boldsymbol{\Sigma} = \mathbf{I}$ and the significance level of $\alpha = 0.05$ when $p = 1000$. Column selection is based on the marginal (or column-wise) variances.

| # selected cols. | Chen et al. (2010) | Fisher (2012) |
|---|---|---|
| All ($p = 1000$) | 0.072 | 0.094 |
| Top 100 | 0.589 | 0.527 |

($n < 20$ and $p > 10000$). For example, the test power is around 0.1 when $n = 15$. Second, even if we select a subset of genes as many practitioners do, the low power of the LC test still makes it difficult to detect differences between two covariance matrices.

#### 2.1 Potential post-selection artifacts

Due to the small sample size, we observe issues with the type-I error rate and power of standard high-dimensional two-sample covariance tests. A naive approach to reduce the number of genes, such as simple data subsetting to conduct covariance test based on differentially expressed genes or gene-wise variance, does not solve the type-I error rate problem from the small sample size $n$, especially with non-Gaussian data. This issue is not limited to two-sample covariance tests; it applies broadly to all types of inferential methods. We provide toy examples to illustrate this problem.

We first consider a one-sample variance testing problem, $H_0 : \Sigma = \mathbf{I}$. We generate simulated data with a sample size of 60 from $N(\mathbf{0}, \mathbf{I})$ where $\mathbf{I}$ is a 1000 by 1000 identity matrix. We compare the type-I error rate of three types of one-sample high-dimensional covariance tests (Chen et al., 2010; Fisher, 2012), using the entirety and the subset of the simulated data. We pick 100 columns of the simulated data with the greatest marginal variances in decreasing order. Table S4 displays the type-I error rate from the analysis. With $\alpha = 0.05$, we observe that the type-I error rates of the tests are not controlled when we use a subset of the simulated data. This demonstrates that subsetting the data does not guarantee the control of the type-I error rate. From this example, we should be keep in mind, when applying the common practice of variance-based filtering for covariance testing, such filtering alone would not resolve the type-I error control issue.

Next, we study whether the covariance test results are affected by the subset selection based on differentially expressed genes for multivariate normal distribution. To this end, we study the type-I error rate of a two-sample covariance test when we only test a subset of the data selected by mean condition. We generate two simulated datasets with $p = 1000$ dimensional multivariate normal data using three covariance structures with 50 columns that have different means, $\boldsymbol{\mu}_1 = (0, \ldots, 0)^{\mathrm{T}} \in \mathbb{R}^{1000}$,

$$\mu_{2,j} = \begin{cases} 5 & j = 1, \ldots, 50 \\ 0 & j = 51, \ldots, 1000 \end{cases}.$$

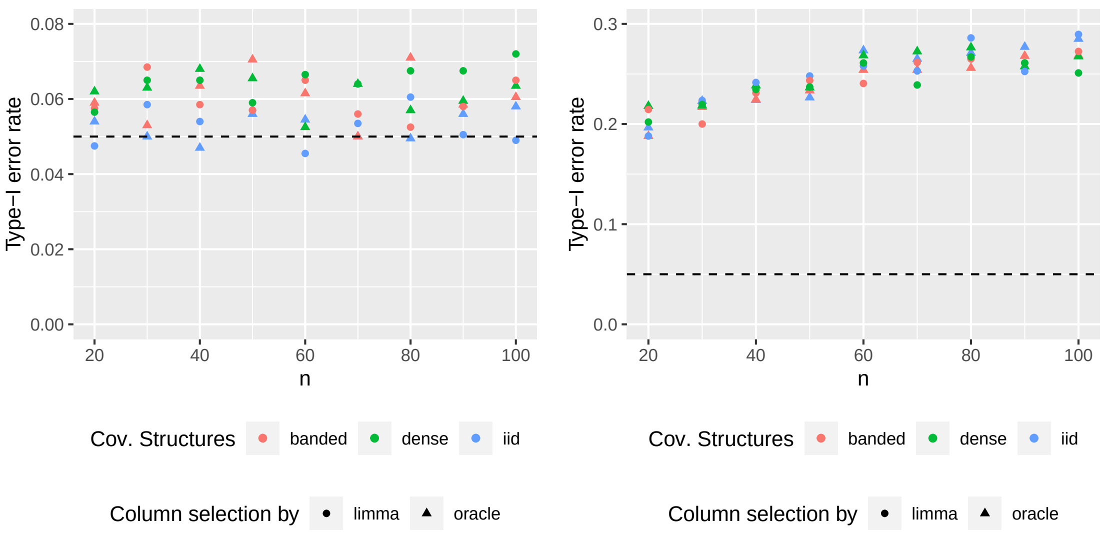


(a) Multivariate normal (b) Multivariate $t$

Figure S3: Type-I error rate of Li and Chen (2012) using selected columns from 1000 dimensional data whose entry-wise means are different (Oracle) and differentially expressed columns by *limma* with a false discovery level 0.05 (DE). Covariance structures in the first row are defined in Section 2. The colors of points indicate covariance or scale matrix structures used in data generation. Shapes of points indicate column selection methods; oracle (○) and *limma* (△). Black-dashed line displays nominal level $\alpha = 0.05$.

Then, we run a two-sample covariance test using the selected columns. Figure S3 displays type-I error rates of the test using subsets of the data with a significance level of $\alpha = 0.05$. We observe that the test controls type-I error rate when we use the subset of the multivariate normal data. This is not surprising since the mean and the covariance are independent under multivariate normal distribution. On the other hand, the test does not control the type-I error rate when we use data from multivariate $t$ distribution for both scenarios. In summary, if there is no guarentee that subsetting genes based on differential mean expression resolve the type-I error rate issue. These two examples suggest that additional methodological adjustments are necessary when the testing procedure fails to control the type I error rate due to small sample size.

### 3 Random projection test

From Section 2 and 2.1, it is clear that the standard high-dimensional covariance tests do not apply to our dataset or its subset because of type-I error rate and power issues. To resolve these issues, we employ the notion of random projections. Wu and Li (2020) considered two-sample covariance testing problems using random projections on one-dimensional space

**Algorithm 1** Random projection algorithm used in the manuscript.

**INPUT:** Two data matrices $\mathbf{X}_1 \in \mathbb{R}^{n_1 \times p}$ and $\mathbf{X}_2 \in \mathbb{R}^{n_2 \times p}$ where both $p$ can be larger than $n_1$ and $n_2$, the number of random projections $K$, the dimension of each random projection $q > 1$, and the significance level of $\alpha$.

**OUTPUT:**

1: Repeat the followings $K$ times.

1. Draw $p$-dimensional vectors $q$ times, $\mathbf{v}'_1, \ldots, \mathbf{v}'_q \in \mathbb{R}^p$, whose entries is from $N(0,1)$, then standardize them $\mathbf{v}_i = \mathbf{v}'_i / \|\mathbf{v}'_i\|_2$. Let $\mathbf{V} = \begin{bmatrix} \mathbf{v}_1 & \cdots & \mathbf{v}_q \end{bmatrix} \in \mathbb{R}^{p \times q}$ be a projection matrix.

2. Compute the projected data matrices $\mathbf{X}'_1 := \mathbf{X}_1 \mathbf{V} \in \mathbb{R}^{n_1 \times q}$ and $\mathbf{X}'_2 := \mathbf{X}_2 \mathbf{V} \in \mathbb{R}^{n_2 \times q}$.

3. Using sub-matrices $\mathbf{X}'_1$ and $\mathbf{X}'_2$, compute the test statistic, $T_i$ for $i = 1, \ldots, K$, from a covariance test (for example, Li and Chen, 2012).

2: Obtaining the most extreme value of $K$ statistics from Step 1 based on the rejection region of the test. For example, if the rejection region of the individual test is $T_i > t$, we use $T = \max_{1 \leqslant i \leqslant K} T_i$, which serves as the test statistic for the proposed test.

3: Comparing the test statistic $T$ with the (simulated) $(1-\alpha) \times 100$-th percentile, denoted by $t_{1-\alpha}$. If $T > t_{1-\alpha}$, reject the null hypothesis.

and the maximum of F-statistics. Instead of testing two large covariance matrices directly, they computed a random projection of the data matrix in $\mathbb{R}^{n \times p}$ to $\mathbb{R}^{n \times 1}$. They used the maximum of $F$-statistics for each random projection, which is from testing the equality of two variances, $\sigma_1^2 = \sigma_2^2$.

We consider a generalized version of Wu and Li (2020) to multi-dimensional space using independently generated random projections. Instead of random projections in one-dimensional space, we consider random projections in multi-dimensional space. The null and alternative hypotheses of the random projection covariance test is the same as those used in two-sample covariance tests:

$$H_0 : \mathbf{\Sigma}_1 = \mathbf{\Sigma}_2, \quad H_a : \mathbf{\Sigma}_1 \neq \mathbf{\Sigma}_2 \tag{3.1}$$

where $\mathbf{\Sigma}_1$ and $\mathbf{\Sigma}_2$ are covariance matrices. The rationale behind the random projection covariance test is that, under the null hypothesis–when the two covariance matrices are equal–the covariances of the projected data should also be similar. The employed testing procedure is summarized in Algorithm 1.

Table S5 displays the simulated critical values for the dimension of projected space $q = 20$ and the maximum of $K \in \{500, 2000\}$ test statistics from Li and Chen (2012) under independent setting defined in Section 2. From the empirical studies, we observe that the simulated critical values from the test statistic are consistent across different dependent

Table S5: Simulated critical values for the dimension of projected space $q = 20$ and the maximum of $K \in \{500, 2000\}$ $\ell_2$-type test statistics under independent setting. The sample sizes of the two groups to be compared are denoted by $n_1$ and $n_2$.

| $K$ | $n_1$ | $n_2$ | 90th | 95th | 97.5th | 98.75th |
|---|---|---|---|---|---|---|
| 500 | 10 | 10 | 2.9961 | 3.0892 | 3.1672 | 3.2123 |
| | 15 | 15 | 3.2905 | 3.4203 | 3.5562 | 3.6837 |
| | 20 | 20 | 3.4841 | 3.6249 | 3.7177 | 3.8971 |
| | 30 | 30 | 3.6690 | 3.8510 | 4.0077 | 4.1631 |
| 2000 | 10 | 10 | 3.1883 | 3.2806 | 3.3604 | 3.4167 |
| | 15 | 15 | 3.5475 | 3.6556 | 3.7776 | 3.8944 |
| | 20 | 20 | 3.7712 | 3.9078 | 4.0351 | 4.1595 |
| | 30 | 30 | 3.9724 | 4.1318 | 4.3120 | 4.4778 |

structures and the dimensions of the original data. We discuss additional details concerning the critical values in Appendix Section A.

We observe that the type-I error rates from the random projection test are controlled even when the sample size is small. Figure S4 displays type-I error rates from the simulated data with 2000 iterations, demonstrating control of type-I error rate across all considered covariance structures. These simulations were conducted with 1000-dimensional simulated data and 500 or 2000 statistics from Li and Chen (2012) with varied dimensions of random projections from 5 to 30, that is, $p = 1000$, $K \in \{500, 2000\}$, and $q \in \{5, 10, 20, 30\}$, using the critical values displayed in Tables S5. Moreover, the type-I error rates are consistent under the considered covariance settings for both values of number of $K$.

Next, we conduct empirical studies on the power of the random projection test. We use the data generation procedure with two different covariance matrices from Section 2. Figure S5 displays the power of the random projection test. We observe that the random projection test has higher power than the standard two-sample covariance test, even when the sample size is less than 20. In other words, if there is a covariance difference between two groups, we can detect the difference using random projection tests when standard tests cannot. Moreover, a larger number of test statistics used to calculate their maximum, larger $K$, leads to higher test power as shown in Figure S5b For this reason, we use the random projection test in Section 4 to resolve the small sample size issue discussed at the beginning of this section.

### 4 Real data analysis

In this section, we provide data analysis results. We conduct two-sample random projection covariance tests in Algorithm 1 using all genes and differentially expressed genes only. This test checks whether the covariance matrices of different groups can be assumed to be equal

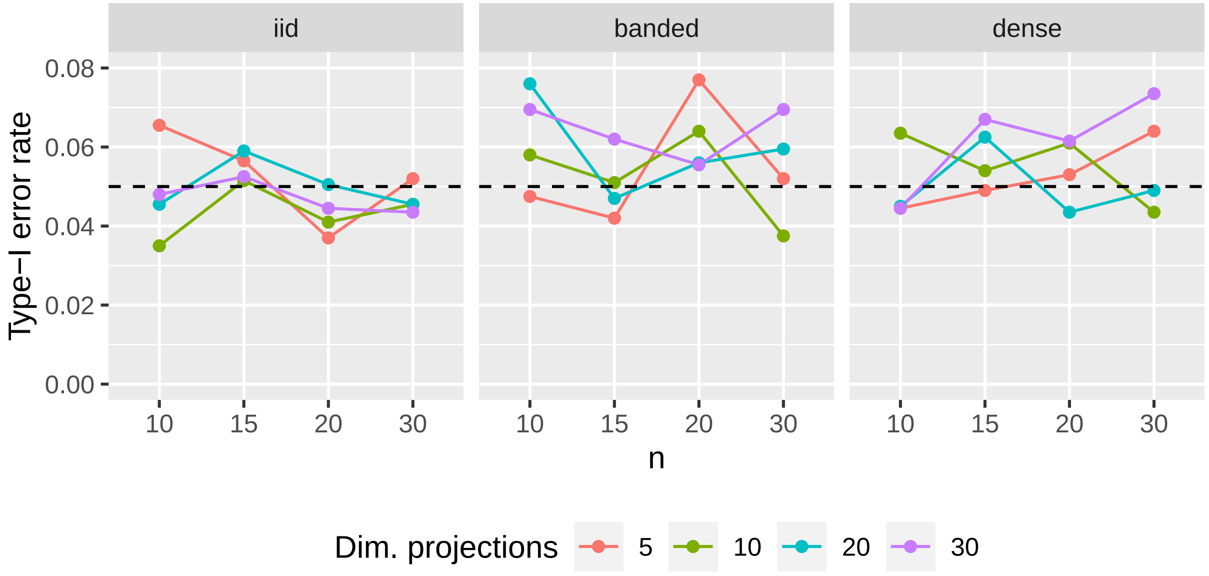


(a) $K = 500$

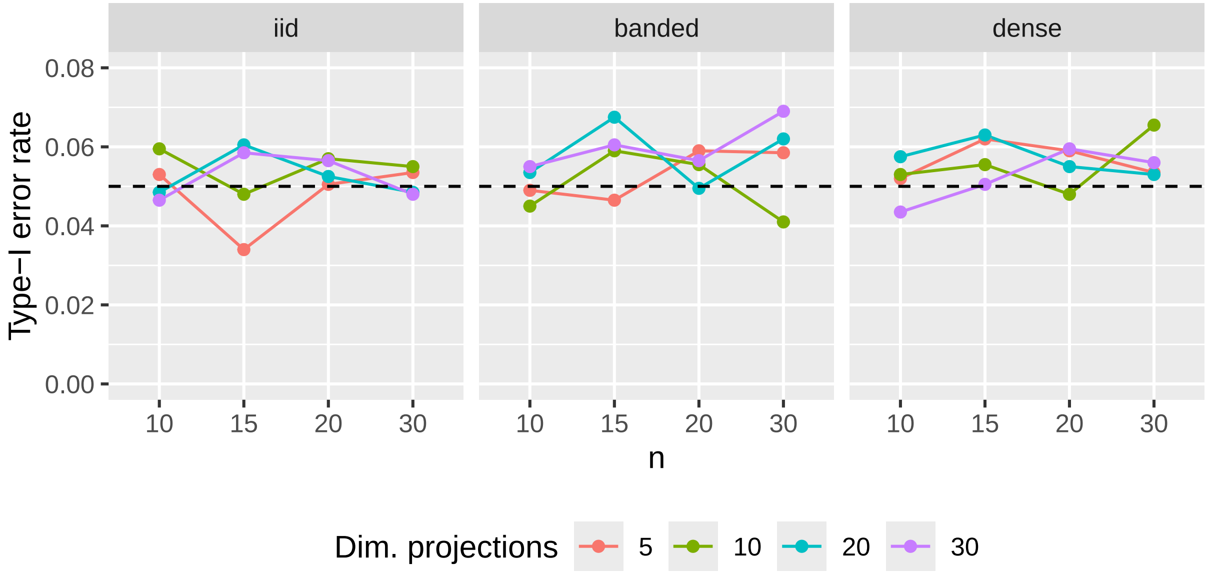


(b) $K = 2000$

Figure S4: Type-I error rate from the random projection test with LC test and the 95th percentiles from Table S5 with various $q \in \{5, 10, 20, 30\}$, the number of random projections, when $p = 1000$ and $K \in \{500, 2000\}$ under the banded matrix structure. The empirical size is computed from 2000 iterations.

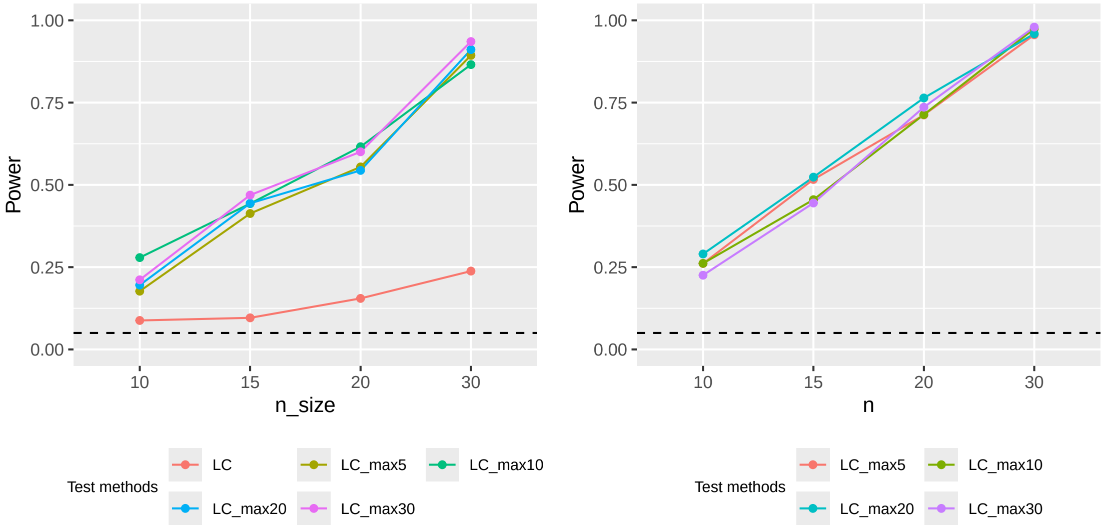


(a) $K = 500$ (b) $K = 2000$

Figure S5: Empirical power of the random projection test with the maximum of $K \in \{500, 2000\}$ LC test statistics with the 95th percentiles from Table S5 with various dimensions of projections $q \in \{5, 10, 20, 30\}$ when $p = 1000$ under the banded covariance matrix setting in (2.1) when $d_j = 1$. The empirical power is computed from 2000 iterations. For comparison, we also report the empirical power of LC test under the same setting from Figure S2.

in the follow-up analysis. Following that, we compare two parts of the correlation networks, DE-DE and NDE-NDE networks, by network summary plots (Maugis et al., 2017) and a network comparison test (Shao et al., 2022). Throughout this section, we use the differential gene expression results from Fischer et al. (2021) to define 'DE' genes as genes differentially expressed based on population effects from generalized linear mixed models.

#### 4.1 Random projection tests

In this subsection, we present results from the random projection test proposed in Section 3 using the residuals from the linear mixed model on the log-transformed gene expression counts. We used 13446 genes for Aripo and 14379 genes for Quare. We exclude one gene in the Aripo dataset from the genes in Table S3 due to 0 group-wise variances for some groups. We aim to assess the statistical significance of covariance differences between pairs of interest: i) LP pred+ versus LP pred-, ii) LP pred+ versus HP pred+, iii) HP pred+ versus HP pred-, iv) LP pred- versus HP pred- where the group labels are defined in Table S1 (see main text for biological justification for comparisons of interest). In other

Table S6: $p$-values from the random projection test using 500 LC test statistics, $K = 500$, where the dimension of random projections per iteration is 20, $q = 20$. We use 13446 genes for Aripo and 14379 genes for Quare.

| | Aripo | Quare |
|---|---|---|
| LP pred+ vs LP pred- | 0.12583 | 0.99433 |
| LP pred+ vs HP pred+ | 0.01490 | 0.00001 |
| HP pred+ vs HP pred- | 1.00000 | 0.06385 |
| LP pred- vs HP pred- | 1.00000 | 0.95684 |

Table S7: Approximated $p$-values from the random projection test on DE genes, 553 in the Aripo dataset and 2909 DE genes in the Quare datasets, using $K = 500$ LC test statistics, where the dimension of the random projection is $q = 20$.

| | Aripo | Quare |
|---|---|---|
| LP pred+ vs LP pred- | 1.00000 | 0.03537 |
| LP pred+ vs HP pred+ | 0.57583 | 0.00000 |
| HP pred+ vs HP pred- | 1.00000 | 0.11046 |
| LP pred- vs HP pred- | 1.00000 | 1.00000 |

words, we consider the following four null hypotheses:

$$
\begin{aligned}
H_{0,LP} &: \boldsymbol{\Sigma}_{LPpred+} = \boldsymbol{\Sigma}_{LPpred-} \\
H_{0,P} &: \boldsymbol{\Sigma}_{LPpred+} = \boldsymbol{\Sigma}_{HPpred+} \\
H_{0,HP} &: \boldsymbol{\Sigma}_{HPpred+} = \boldsymbol{\Sigma}_{HPpred-} \\
H_{0,NP} &: \boldsymbol{\Sigma}_{LPpred-} = \boldsymbol{\Sigma}_{HPpred-}
\end{aligned}
$$

We employ the proposed test with the maximum of statistics from Li and Chen (2012), using 500 random projections in a 20-dimensional space ($q = 20$ and $K = 500$). To obtain $p$-values, we simulate random projection test statistics from 100000 repetitions under the *i.i.d.* setting when $p = 1000$, $q = 20$, and $K = 500$. We used *i.i.d.* setting based on the empirical studies in Section A, which demonstrate the simulated critical values have consistent trends across various covariance setups and the dimension of the data, $p$. Table S6 displays the approximated $p$-values from the random projection test. The random projection test results imply covariance differences across groups in the Aripo and Quare datasets with the nominal family-wise error rate of 0.1 with Bonferroni correction. The test rejects $H_{0,P}$ at a significance level of 0.025 for both datasets.

#### 4.2 Two-sample covariance tests with DE genes

We also compare sub-matrices of the entire covariance matrices using the random projection test. We focus on the covariance of DE genes:

$$\begin{aligned}
H_{0,LP,DE} :& \boldsymbol{\Sigma}_{LP\ pred+,DE} = \boldsymbol{\Sigma}_{LP\ pred-,DE} \\
H_{0,P,DE} :& \boldsymbol{\Sigma}_{LP\ pred+,DE} = \boldsymbol{\Sigma}_{HP\ pred+,DE} \\
H_{0,HP,DE} :& \boldsymbol{\Sigma}_{HP\ pred+,DE} = \boldsymbol{\Sigma}_{HP\ pred-,DE} \\
H_{0,NP,DE} :& \boldsymbol{\Sigma}_{LP\ pred-,DE} = \boldsymbol{\Sigma}_{HP\ pred-,DE}.
\end{aligned}$$

Table S7 displays $p$-values of random projection tests. Using a family-wise error rate of 0.1 with Bonferroni correction, we conclude that the covariance difference between LP pred+ and HP pred+ groups for Quare data set is statistically significant. On the other hand, we do not have enough evidence to conclude that there are covariance differences for any pairs in the Aripo dataset.

#### 4.3 Correlation network comparison

In addition to the covariance matrix comparisons, we conduct correlation network comparisons between sets of DE and NDE genes. We clarify two things: (i) how to define *correlation network*; (ii) how to compare two networks with different sizes. To construct correlation networks for each group in Table S1, we employ high-dimensional correlation tests with the null hypotheses $H_{0ij} : \rho_{ij} = 0$ for $1 \leqslant i < j \leqslant p$ by Cai and Liu (2016), using the nominal false discovery rate (FDR) level of 0.05. Specifically, we test the correlation of residuals from linear mixed models to determine whether there is a connection between two genes, that is, an edge in the network. As a result, we obtained adjacency matrices of undirected simple graphs. For the second question, we employ both exploratory analysis and statistical inference. The former is conducted by network summary plots (Maugis et al., 2017). For the latter, we run a two-sample network test proposed by Shao et al. (2022).

The main idea of network summary plots is subsampling nodes and counting cycles of the sampled subgraphs. The shape of the summary plots differs if the two network structures are different. For example, if a network is a tree, then there are no cycles in the network. Therefore, only the first component of the summary plot, trees with three nodes also known as v-shape, displays positive values. On the other hand, for a complete graph, all the components in the summary plot have the maximum values.

It is expected to have an overall inflation of edge densities and subgraph densities such as triangle density due to a greater number of rejections as the level of false discovery rate (FDR) increases. Figure S6 shows the DE genes network summary plots for the HP pred+ group from the Aripo drainage by varying FDR levels, $\alpha \in \{0.01, 0.05, 0.1\}$. Nevertheless, the order of y-axis values in the summary plots remains the same, which implies

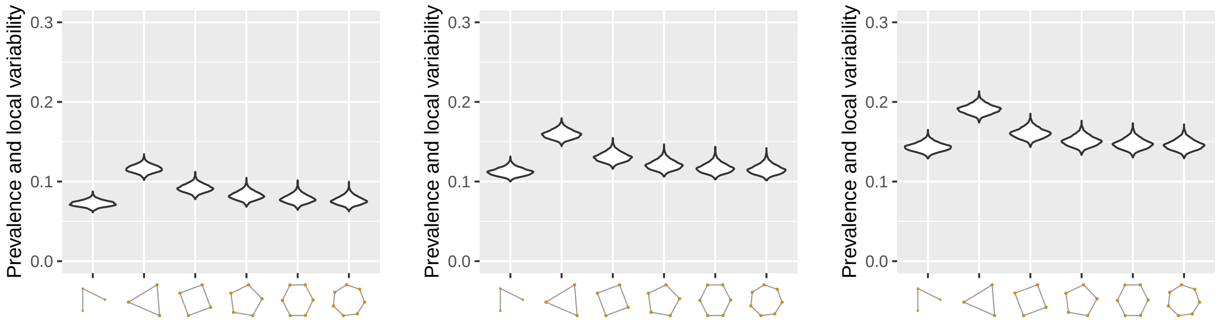


(a) $\alpha = 0.01$ (b) $\alpha = 0.05$ (c) $\alpha = 0.1$

Figure S6: Network summary plots for correlation networks of Aripo HP pred+ DE genes constructed by Cai and Liu (2016) with different FDR level, $\alpha \in \{0.01, 0.05, 0.1\}$.

the corresponding network structure remains the same even if the sparsity is different. We observe similar phenomena across all cases. Hence, we focus on the correlation networks from $\alpha = 0.05$ for the rest of this section.

Table S8 provides the summary statistics of the constructed correlation networks by Cai and Liu (2016) with a false discovery level of 0.05. Edge densities of networks from the Aripo dataset are in the 10–13% range, those from the Quare dataset are widespread from 6.3% to 34%. Positive values of degree assortativity imply that nodes with similar degrees are likely linked. Hubs in the networks are linked, while they do not link to nodes with small degrees. On the other hand, small degree nodes are connected. The global clustering coefficient, defined by the ratio of triangles over connected triplets, varies from 0.34 to 0.41 in the Aripo dataset and from 0.28 to 0.76 in the Quare dataset.

Next, we compare the network summary plots of the constructed correlation networks of interest. Figures S7 and S8 display the network summaries for the Aripo and Quare datasets, respectively. We do not observe visual differences between DE and NDE in Aripo HP pred+ and Aripo HP pred-. For all other cases, NDE networks have larger summary values than DE networks.

Lastly, we employ a network comparison test proposed by Shao et al. (2022), which can test two networks with different sizes via the difference in sparsity-adjusted density of a subgraph, defined as the subgraph density divided by the edge densities. If two networks share a structure, the difference should be small. For the following test procedure, we use v-shapes, triangles, and 3-stars, displayed in Figure S9. These three subgraphs, a smaller graph in a larger graph, have conceptual attractiveness and computational advantages due to their simple structure, as they provide insights into connectivity and community prevalence while requiring less computational time than other subgraphs.

Table S9 displays $p$-values of the test comparing DE correlation network with NDE correlation networks using v-shapes, triangles, and 3-stars. The test was conducted with a two-sided alternative hypothesis, assessing whether the two networks have different sparsity-

Table S8: Summary statistics for correlation networks used in the data analysis. The number of nodes is denoted by $|V|$. We denote the edge density of a network by $\rho_n = |V|/\binom{n}{2}$, degree assortativity (Newman, 2002) by $r_d$, and the global clustering coefficient (Barabási, 2017) by $c_\Delta$.

| Aripo | $\rho_n$ | $r_d$ | $c_\Delta$ |
|---|---|---|---|
| HP pred+, DE | 0.1073 | 0.3398 | 0.3382 |
| HP pred+, NDE | 0.1120 | 0.3659 | 0.3534 |
| HP pred-, DE | 0.0976 | 0.3964 | 0.3403 |
| HP pred-, NDE | 0.1076 | 0.4313 | 0.3557 |
| LP pred+, DE | 0.1035 | 0.3254 | 0.3255 |
| LP pred+, NDE | 0.1122 | 0.4246 | 0.3528 |
| LP pred-, DE | 0.1136 | 0.4425 | 0.3736 |
| LP pred-, NDE | 0.1272 | 0.4866 | 0.4115 |
| Quare | $\rho_n$ | $r_d$ | $c_\Delta$ |
| HP pred+, DE | 0.0633 | 0.4119 | 0.2762 |
| HP pred+, NDE | 0.0807 | 0.4320 | 0.3490 |
| HP pred-, DE | 0.1207 | 0.4380 | 0.3933 |
| HP pred-, NDE | 0.1907 | 0.3824 | 0.5362 |
| LP pred+, DE | 0.1220 | 0.4385 | 0.5035 |
| LP pred+, NDE | 0.2388 | 0.3037 | 0.6675 |
| LP pred-, DE | 0.1944 | 0.3581 | 0.6422 |
| LP pred-, NDE | 0.3403 | 0.2172 | 0.7557 |

Table S9: $p$-values from the two-sample network test with a two-sided alternative (Shao et al., 2022) using v-shapes, triangles, and 3-stars on correlation networks of population DE and NDE genes.

| | Subgraph | HP pred+ | HP pred- | LP pred+ | LP pred- |
|---|---|---|---|---|---|
| Aripo | v-shape | 0.1344 | 0.0106 | $< 0.0001$ | 0.0112 |
| | triangle | 0.4620 | 0.2834 | 0.0310 | 0.3423 |
| | 3-star | 0.9996 | 0.7589 | 0.1529 | 0.5425 |
| Quare | v-shape | $< 0.0001$ | $< 0.0001$ | 0.9751 | $< 0.0001$ |
| | triangle | $< 0.0001$ | 0.0959 | $< 0.0001$ | $< 0.0001$ |
| | 3-star | 0.1322 | $< 0.0001$ | $< 0.0001$ | $< 0.0001$ |

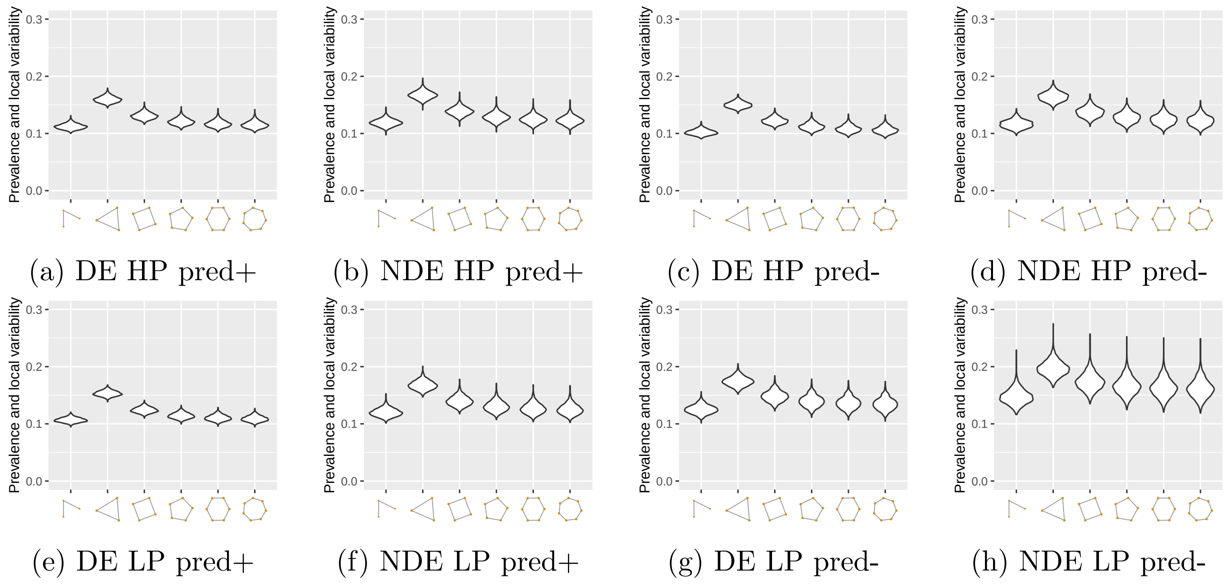


(a) DE HP pred+ (b) NDE HP pred+ (c) DE HP pred- (d) NDE HP pred-

(e) DE LP pred+ (f) NDE LP pred+ (g) DE LP pred- (h) NDE LP pred-

Figure S7: Network summary plots of correlation networks for Aripo dataset.

adjusted subgraph densities, defined as subgraph densities divided by network sparsity. At a significance level of 0.05, we conclude a statistically significant difference between DE and NDE networks for all groups from Quare drainage. On the other hand, In the Aripo drainage, we identify statistically significant differences between DE and NDE in all groups except HP pred+. Other comparisons, such as one-sided tests, are presented in the manuscript.

We also perform the test with one-sided alternatives. In addition to the one-sided test results presented in the manuscript, we provide an additional comparison here. Table S10 presents $p$-values from the "less" alternative indicating the DE network has a smaller sparsity-adjusted subgraph density than NDE networks in the specified setting, while "greater" implies the opposite. The test determines the direction in network comparisons, identifying whether one network has a higher sparsity-adjusted density than the other. For example, in the Aripo drainage, the DE networks have less sparsity-adjusted v-shape densities than the NDE networks in the HP pred-, LP pred+, and LP pred- groups. Moreover, in Quare drainage, the DE has higher v-shape and triangle sparsity-adjusted densities than the NDE in HP pred+ group, whereas the DE networks in LP pred+ and LP pred- have lower triangle and 3-star sparsity-adjusted densities than their NDE counterparts.

##### 4.3.1 Simulation studies on the validity

We check that the test is valid for a very unbalanced scenario since DE and NDE networks have different numbers of vertices. We apply the test on two generated networks by the network model used in Bickel and Chen (2009) with the number of vertices, $n_1, n_2 \in$

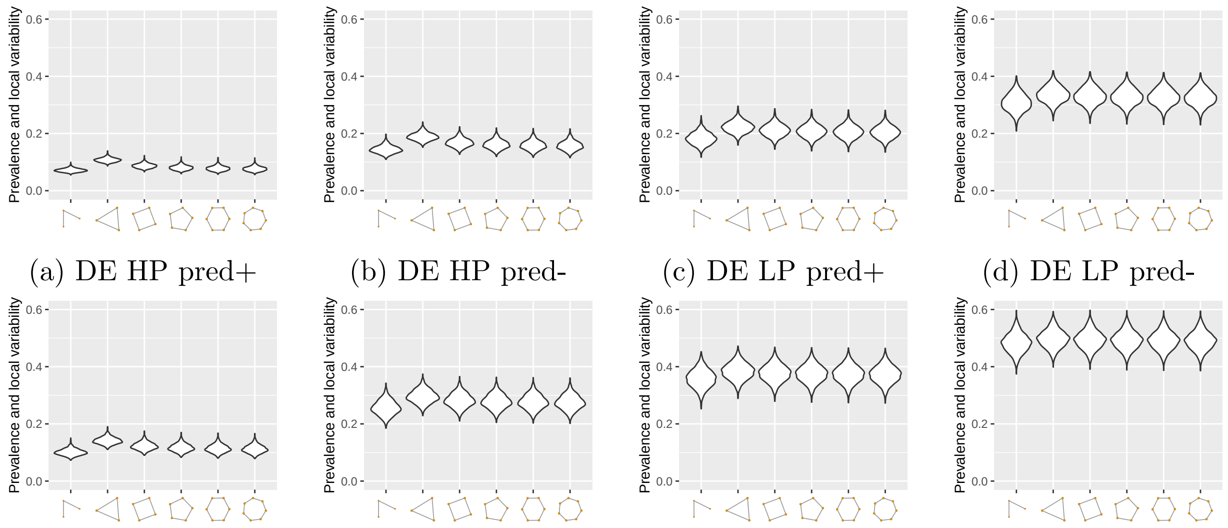


(a) DE HP pred+ (b) DE HP pred- (c) DE LP pred+ (d) DE LP pred-

(e) NDE HP pred+ (f) NDE HP pred- (g) NDE LP pred+ (h) NDE LP pred-

Figure S8: Network summary plots of correlation networks for Quare dataset.

Table S10: Network comparisons of DE vs NDE genes within treatment groups and across datasets. Comparisons of sparsity-adjusted subgraph densities tested the alternatives that DE gene networks had smaller or larger subgraph density than NDE networks. P-values from one-sided alternative tests (Shao et al., 2022) are reported for the v-shape, triangle, and 3-star subgraph types. “DE $<$ NDE” implies that the DE network has a smaller sparsity-adjusted subgraph density than the NDE network in the specified setting, while “DE $>$ NDE” implies the opposite.

| | ARIPO drainage | | | QUARE drainage | | |
|---|---|---|---|---|---|---|
| | v-shape | triangle | 3-star | v-shape | triangle | 3-star |
| DE <NDE | | | | | | |
| HP pred- | 0.0051 | 0.8611 | 0.6149 | <0.0001 | 0.9523 | 1.0000 |
| HP pred+ | 0.0676 | 0.2330 | 0.5027 | <0.0001 | <0.0001 | 0.0665 |
| LP pred - | 0.0056 | 0.1700 | 0.2650 | 1.0000 | 1.0000 | 1.0000 |
| LP pred+ | <0.0001 | 0.0152 | 0.0749 | 0.4844 | 1.0000 | 1.0000 |
| DE >NDE | | | | | | |
| HP pred- | 0.9948 | 0.1389 | 0.3742 | 1.0000 | 0.0482 | <0.0001 |
| HP pred+ | 0.9358 | 0.7676 | 0.5013 | 1.0000 | 1.0000 | 0.9347 |
| LP pred - | 0.9938 | 0.8277 | 0.7304 | <0.0001 | <0.0001 | <0.0001 |
| LP pred+ | 1.0000 | 0.9850 | 0.9252 | 0.5122 | <0.0001 | <0.0001 |

$\{20, 40, 80, 160, 320\}$, from a function $f(x, y) = (1.71/8) \cdot (x + y)^2 \mathbb{I}(x \neq y)$. Specifically, we draw $n_1$ random samples from the uniform distribution from 0 and 1, denoted by $\{\xi_i\}_{i=1}^{n_1}$. Following that, we generate Bernoulli samples from $A_{ij} \sim Bern\{f(\xi_i, \xi_j)\}$, and $A_{ji} = A_{ij}$. If $A_{ij} = 1$, then there is an edge between vertices $i$ and $j$. We repeat this generating proce-

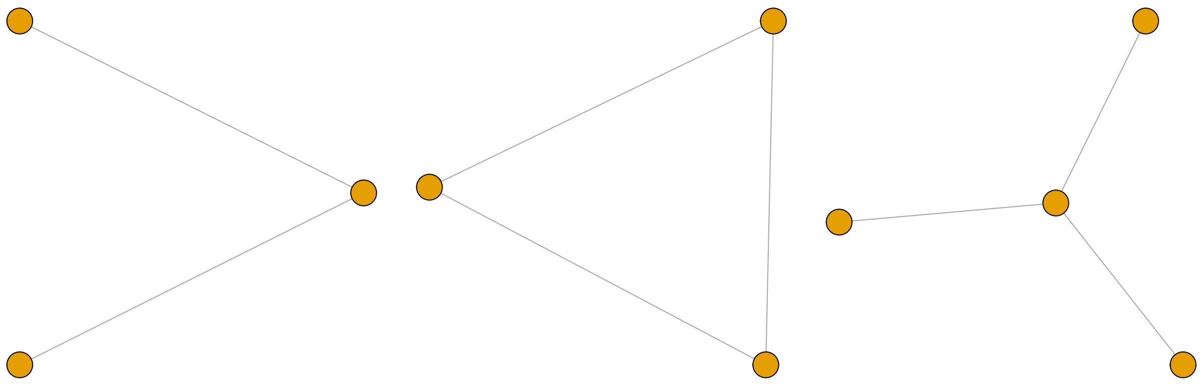

(a) v-shape (b) triangle (c) 3-star

Figure S9: List of sub-graphs used in two-sample network test for Tables 3 and S9.

Table S11: Type-I error rates of two-sample network test (Shao et al., 2022) using triangles with the significance level of $\alpha = 0.1$ by varying $(n_1, n_2)$. Networks are generated from a function $f(x, y) = (1.71/8) \cdot (x + y)^2 \mathbb{I}(x \neq y)$. Type-I error rate is computed from 200 repetitions.

| $n_1 \backslash n_2$ | 20 | 40 | 80 | 160 | 320 |
|---|---|---|---|---|---|
| 20 | 0.130 | 0.090 | 0.150 | 0.120 | 0.145 |
| 40 | 0.090 | 0.090 | 0.110 | 0.130 | 0.135 |
| 80 | 0.110 | 0.075 | 0.095 | 0.105 | 0.105 |
| 160 | 0.145 | 0.080 | 0.070 | 0.080 | 0.055 |
| 320 | 0.135 | 0.065 | 0.105 | 0.085 | 0.110 |

dure for the other network with $n_2$ nodes, then run the test proposed by Shao et al. (2022) using triangles. Then, we repeat this procedure 200 times for each combination of $(n_1, n_2)$. Table S11 displays the type-I error rates of the test with the nominal level of $\alpha = 0.1$. We observe that the test controls the type-I error rate, with the values in the table remaining close to 0.1, even when the two sample sizes are different. Hence, we can use the test for our study since we are interested in comparing DE and NDE networks, which have very different network sizes.

### References

Barabási, A.-L. (2017). *Network Science* (4th print. ed.). Cambridge: Cambridge University Press.

Bickel, P. J. and A. Chen (2009, December). A nonparametric view of network models and Newman–Girvan and other modularities. *Proceedings of the National Academy* *of Sciences 106*(50), 21068–21073. http://www.pnas.org/lookup/doi/10.1073/pnas.0907096106.

Cai, T., W. Liu, and Y. Xia (2013). Two-Sample Covariance Matrix Testing and Support Recovery in High-Dimensional and Sparse Settings. *Journal of the American Statistical Association 108*(501), 265–277.

Cai, T. T. and W. Liu (2016). Large-scale multiple testing of correlations. *Journal of the American Statistical Association 111*(513), 229–240. PMID: 27284211.

Chen, S. X., L.-X. Zhang, and P.-S. Zhong (2010). Tests for High-Dimensional Covariance Matrices. *Journal of the American Statistical Association 105*(490), 810–819.

Fischer, E. K., Y. Song, K. A. Hughes, W. Zhou, and K. L. Hoke (2021). Nonparallel transcriptional divergence during parallel adaptation. *Molecular Ecology 30*(6), 1516–1530.

Fisher, T. J. (2012). On testing for an identity covariance matrix when the dimensionality equals or exceeds the sample size. *Journal of Statistical Planning and Inference 142*(1), 312–326.

Li, J. and S. X. Chen (2012). Two sample tests for high-dimensional covariance matrices. *The Annals of Statistics 40*(2), 908–940.

Maugis, P.-A. G., S. C. Olhede, and P. J. Wolfe (2017). Topology reveals universal features for network comparison. *arXiv:1705.05677 [cs, math, stat]*.

Newman, M. E. J. (2002). Assortative mixing in networks. *Physical Review Letters 89*, 208701.

Shao, M., D. Xia, Y. Zhang, Q. Wu, and S. Chen (2022). Higher-order accurate two-sample network inference and network hashing. arXiv:2208.07573 [math, stat].

Wu, T.-L. and P. Li (2020). Projected tests for high-dimensional covariance matrices. *Journal of Statistical Planning and Inference 207*, 73–85.

Yu, X., D. Li, and L. Xue (2024). Fisher's Combined Probability Test for High-Dimensional Covariance Matrices. *Journal of the American Statistical Association 119*(545), 511–524.

### A Random projection test with LC test statistics

In this section, we provide extra simulation results for critical values of the random projection test with the maximum of $L_2$ type test statistics. This includes the effects of the sample size ($n$), data dimension ($p$), and covariance structures on the critical values.

We obtain the 95th percentile from $B = 2000$ replications, which is obtained by taking the maximum of $K = 500$ test statistics from Li and Chen (2012). We use the number of random projections $q = \{5, 10, 20, 30\}$. For simplicity, we generate two data matrices $\mathbf{X}_1$ and $\mathbf{X}_2$ from a common distribution $N(\mathbf{0}, \mathbf{\Sigma})$ with $(n_1, n_2) \in \{(10, 10), (15, 15), (20, 20), (30, 30)\}$. We take $n_1 + n_2$ random samples from $N(\mathbf{0}, \mathbf{\Sigma})$ where $\mathbf{\Sigma} = \mathbf{D}\widetilde{\mathbf{\Sigma}}\mathbf{D}$, $\mathbf{D} = \text{diag}(d_1, \ldots, d_p)$, $d_j \sim \text{uniform}(1, 5)$, and $\widetilde{\mathbf{\Sigma}} = \{\widetilde{\sigma}_{j,k}\}_{1 \leqslant j,k \leqslant p}$ is defined as follows:

1. (independent) $\widetilde{\mathbf{\Sigma}} = \mathbf{I}$.
2. (Banded) $\widetilde{\sigma}_{j,j} = 1$, $\widetilde{\sigma}_{j,j\pm1} = 0.6$, $\widetilde{\sigma}_{j,j\pm2} = 0.3$, and $\widetilde{\sigma}_{j,k} = 0$ when $|j - k| \geqslant 3$.
3. (Dense) $\widetilde{\sigma}_{j,k} = 0.6^{|j-k|}$.

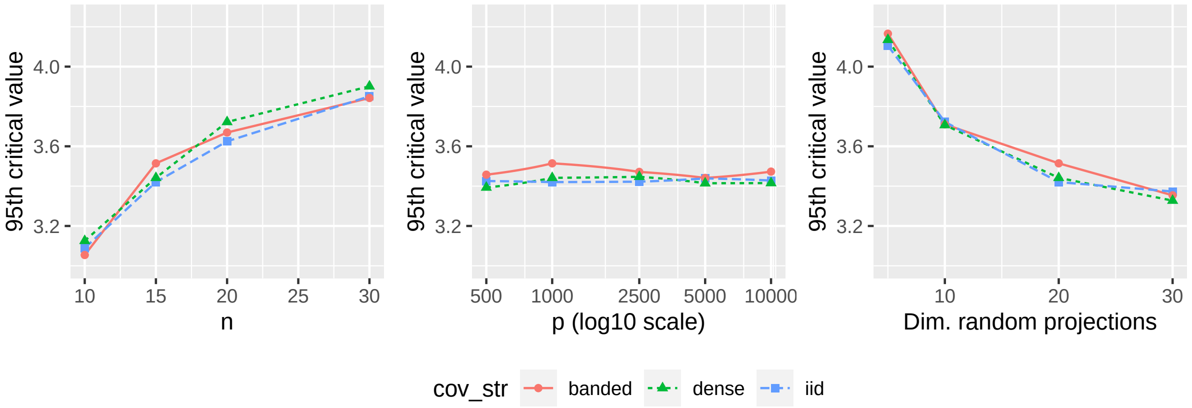


Figure S10: Critical values by the sample size (left), dimension of the data (middle), and dimension of random projections (right). Each line type indicates types of covariance structures.

Overall, we observe the values from the considered covariance structures are similar for each setting. On the other hand, the values differ based on sample size (denoted by $n$), dimension of the data (denoted by $p$), and dimension of random projections (denoted by $q$). Figure S10 displays the simulated 95th percentiles by $n$, $p$, and $q$ with 500 random projections ($K = 500$) from 2000 repetitions. The left figure displays the critical values when $p = 1000$ and $q = 20$. As $n$ increases, the critical values increase too. The middle figure displays the critical values when $n = 15$ and $q = 20$. The values are consistent with different $p$. The right figure displays the critical values with respect to the projection dimension $q$ when $n = 15$ and $p = 1000$. We observe smaller critical values as the dimension of projection increases. Hence, we use the critical values from the independent setting with $(n_1, n_2) \in \{(10, 10), (15, 15)\}$ and $p = 1000$ in Section 4.
